# Kinetic Control of Out-Of-Equilibrium Dynamics in the RhoA Signaling Cascade Shapes Actomyosin Contractility

**DOI:** 10.64898/2026.02.08.704688

**Authors:** Serena Prigent Garcia, Étienne Pinard, Camille N. Plancke, Jing Li, Shashi Kumar Suman, Loan Bourdon, Christelle Gally, Taeyoon Kim, François B. Robin

## Abstract

Cellular functions rely on the precise timing of signal transmission through sequential activation cascades, yet the origin and functional role of signaling delays remain poorly understood. Here, we dissect the temporal organization of the RhoA signaling cascade during pulsed actomyosin contractility in the early C. elegans embryo. We uncover a stereotypical delay between upstream RhoA/ROCK activation and downstream myosin II recruitment. Using TIRF single-molecule microscopy, we show that this delay arises from binding and unbinding kinetics of myosin rather than from slow biochemical reactions. A simple and versatile kinetic model parameterized by these measurements accurately predicts the temporal evolution of myosin accumulation and reveals active control of the dynamic range of the cascade. Perturbing actin and myosin turnover experimentally confirms these predictions, and numerical simulations show that the delay between actin and myosin plays a critical role in force deployment during pulsed contraction. Together, our results indicate that kinetic delays in signaling cascades are not simply tolerated during morphogenesis, but actively shape force deployment in the cell.

## Introduction

Signaling cascades are a widespread biological strategy to respond to, propagate and shape extracellular and intracellular inputs. Operating across multiple spatial and temporal scales over a wide repertoire of molecular substrates –from Rho GTPases, to second messengers, to transcriptional regulation– signaling cascades carry critical information within biological systems (Novák and Tyson 2008; Kholodenko 2006; Ferrell 2013; Goldbeter 1996). The architecture and tuning of these cascades underlie a variety of behaviors, ranging from signal magnitude or sensitivity amplification (Jr., Goldbeter, and Stock 1982), ultrasensitivity (Huang and Ferrell 1996) or bi-stability (Markevich, Hoek, and Kholodenko 2004; Mori, Jilkine, and Edelstein-Keshet 2008), to cyclic oscillations (Pomerening, Sontag, and Ferrell 2003) or chaotic dynamics (Gérard and Goldbeter 2012; Yamaguchi, Ode, and Ueda 2021), and even intricate dynamics within seemingly simple 3-component networks (Ma et al. 2009). While mathematical models provide a glimpse on how these signaling cascades are tuned, and even as optogenetic tools offer a handle to control cascade activation while monitoring its propagation (Valon *et al*, 2015, de Seze *et al*, 2025), experimental measurements on dynamic systems still remain notably constrained. The experimental dissection of the molecular mechanisms underpinning the spatial and temporal dynamics of these activation cascades therefore represent a key objective to understand how signals propagate in the cell.

To dissect the dynamics of hierarchical signaling cascades, we decided to focus on the RhoA cascade, which orchestrates actomyosin recruitment during pulsed contractions. Small RhoGTPases represent a key example of signaling cascades, yet the dynamics governing the activation of the downstream players within these cascades remains largely uncharted. Over the past 25 years, RhoA in particular has emerged as a central regulator of morphogenesis (Hariharan et al. 1995; Eaton et al. 1995), from *Drosophila* gastrulation, germband extension and amnioserosa contraction during dorsal closure (Barrett, Leptin, and Settleman 1997; Munjal et al. 2015; Solon et al. 2009), cells convergent extension in *Xenopus* early embryo (H. Y. Kim and Davidson 2011), to compaction in the early mouse embryo (Maître et al. 2015). In *C. elegans*, RhoA contributes to diverse processes including spermatheca contraction (Tan and Zaidel-Bar 2015), epidermal cell migration (Wallace et al. 2018), embryonic elongation (Diogon et al. 2007; Gally et al. 2009), and plays a critical role in embryo polarization (Motegi and Sugimoto 2006; Schonegg et al. 2007). At the molecular level, RhoA drives the dynamic remodeling of cortical actomyosin networks and actomyosin contractility through a dual effect on F-actin and Myosin II. RhoA activates and recruits the *mDia*/*Dia* formin homolog CYK-1, which processively elongates actin filaments (Kovar and Pollard 2004; Costache et al. 2022). RhoA also activates and recruits the Rho kinase/ROCK (LET-502), which phosphorylates Myosin Regulatory Light Chain and inhibits Myosin Phosphatase (MEL-11) (Diogon et al. 2007; Piekny and Mains 2002; Wissmann et al. 1997; Wissmann, Ingles, and Mains 1999) resulting ultimately in Myosin II activation and cortical recruitment (Fig. 1A).

**Figure 1.**
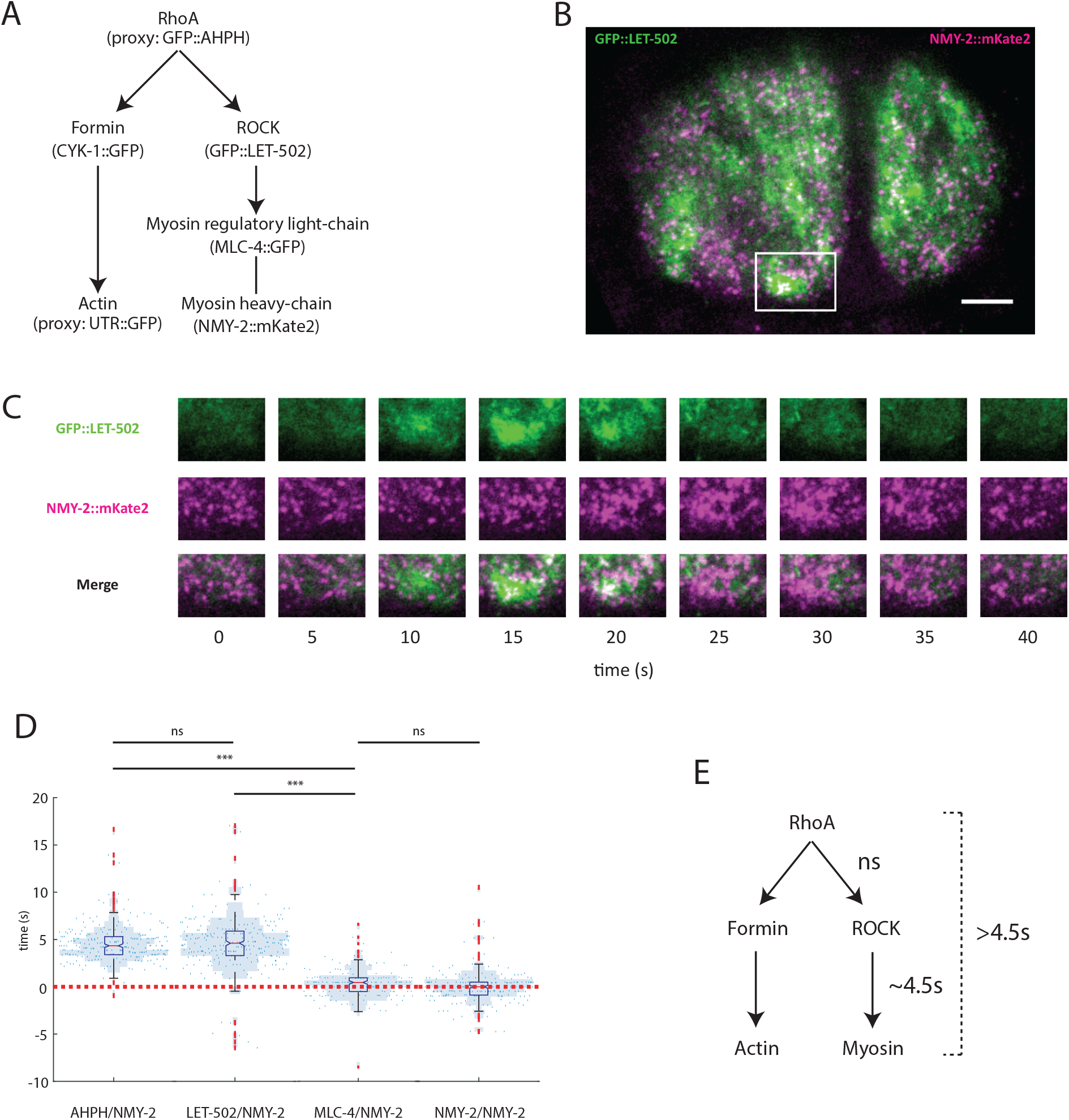
RhoA hierarchical signaling cascade reveals a delay between RhoA, ROCK and Myosin. **(A)** RhoA activation cascade of Actomyosin pulsed contractions with the different strains and proxys used in (D). **(B)** Near-TIRF microscopy image of a 2-cell stage *C. elegans* embryo showing ROCK in green (GFP::LET-502) and Myosin in magenta (NMY-2::mKate2), scale bar: 5 μm. **(C)** Timelapse of the pulsed contraction in white box in (A). **(D)** Quantifications of time delay. Blue dots: time delay distribution red over green channel, red dotted line: time delay of 0 s, blue shade: histogram of distribution. N(embryos) ≥ 10, N(pulses) ≥ 165 (See Supplementary Table S1 and S2 for statistical details), ns: not significant, ***: p-value ≤ 0.001. **(E)** Summary of the observed time delay within the RhoA activation cascade.

Dynamically, RhoA local activation drives actomyosin activation leading to continuous contraction of actomyosin cables, but also frequently occurs in pulses (Miao & Blankenship 2020). Pulsed actomyosin contractility represents a widespread mode of actomyosin contractility, and drives a broad variety of morphogenetic processes, from cell and tissue invagination during gastrulation to cell behavior coordination. Depending on the biological context, the period of pulsed contractions varies widely, from 30 s in *C. elegans* (Michaux et al. 2018), to 2 min in *Drosophila* gastrulation or germband extension (Martin, Kaschube, and Wieschaus 2009; Munjal et al. 2015), to 6 min in *Drosophila* follicle cells (He et al. 2010). While recent work has started to explore how the duration and iteration of contractions is linked to deformation reversibility and effectiveness (Cavanaugh et al. 2020; Staddon et al. 2019), the molecular mechanisms underpinning signal transduction in this cascade still remain unclear, in particular regarding the delay between steps of the cascade (Michaux et al. 2018), the molecular origins of these delays, and their impact on cellular mechanics.

Here, we first describe the time delay between consecutive steps of this hierarchical cascade. Using single-molecule microscopy, we provide dynamic measurement in live embryos of the binding rate (number of molecules binding to the cortex per time unit, noted *k*_*app*_) and the unbinding rate (fraction of molecules unbinding from the cortex per time unit, noted *k*_*off*_), thereby exposing dynamic modulations of the signaling kinetics. Using a simple mathematical model, we then explore numerically the impact of these modulations. Based on this simple model, we derive the *signaled concentration* –a concentration that the system would reach if the binding and unbinding rates were to remain constant– which monitors the out-of-equilibrium dynamics of the system. We subsequently use our dynamic measurements of *k*_*app*_ and *k*_*off*_ to infer the *signaled concentration in vivo*. To modulate pulse period and challenge our model, we then use genetic perturbations to modulate F-actin assembly/disassembly and Myosin II binding/unbinding rates. Finally, we leverage computer simulations to explore the impact of the delay on F-actin contraction. Together, our results show that kinetic delays are an integral part of signaling cascades and strongly impact mechanical outputs, affecting the effective deployment of actomyosin contractility.

## Results

### Pulsed contractions as model to explore signal propagation in a signaling cascade

The RhoA cascade display four key features that were of interest to us. First, it is highly hierarchical, with two branches derived from the same upstream signal, a *F-actin* branch and a *Myosin II* branch. Second, these two branches of the cascade contribute to generate the output dynamics, raising the question of the importance and modalities of the synchronization between two outcomes in parallel transduction pathways from a single input. Third, the cascade displays a pulsatile activation, which we effectively used as iterative cycles of activation to study signal propagation in the cascade. Finally, the RhoA signaling cascade is interfaced with a mechanical output, raising generic questions regarding how mechanical and biochemical signaling can intertwine.

In the zygote, however, pulsed contractions are associated with deep invaginations of the cortex, potentially affecting measurements of pulse kinetics, hampering a detailed analysis of pulse kinetics, in particular for single-molecule microscopy analysis. We therefore decided to focus on the dynamics of the cascade at the 2-cell stage. We have previously shown that RhoA-driven pulsed activation causes local cortical actomyosin contractility (Michaux et al. 2018), potentially affecting the tension distributed in the actomyosin cortex and causing mechanosensitive recruitment of cascade components. The amount of contraction itself, however, remains reasonably small (∼5%), arguably affecting only marginally the local concentration of the components of the cascade. Finally, the RhoA signaling cascade unfolds at the cell surface, greatly simplifying the observation.

### Myosin II accumulation is delayed compared to RhoA activation

We first decided to proceed to an in-depth description of the temporal kinetics of the RhoA signaling cascade. We deployed a general strategy to measure the dynamics of accumulation of the sequential players of the activation cascade, focusing initially on the Myosin II branch. Using strains co-expressing Myosin Heavy-Chain NMY-2 labeled with mKate2 (NMY-2::mKate2, (Dickinson et al. 2017)) along with GFP-tagged probes for each individual player of the cascade, we observed the cell cortex with near-total internal reflection (TIRF) microscopy to assess Myosin II accumulation and pulse initiation (Fig. 1A). We visually identified pulses as local accumulations of Myosin II, spanning several microns, in the anterior (AB) cells during interphase at the 2-cell stage, then quantified and compared the accumulation of Myosin II with the accumulation of the GFP-tagged probe (Fig. 1B, C).

To establish a baseline control for our measurements, we used a strain co-expressing on the one hand NMY-2 fused with mKate2 (red fluorescent protein) at the genomic locus, and on the other hand an ectopic copy of NMY-2 fused with GFP inserted in the genome (Nance 2003). Simple observation revealed that both fusion proteins were expressed in overlapping patterns, with almost identical spatial distributions around the center of the pulse (Movie S1). We then measured the accumulation kinetics of NMY-2::GFP and NMY-2::mKate2. As expected, we observed on average no significant overall difference in (Fig. 1D, Fig. S1), confirming the validity of our approach. Interestingly, we observed that most delays were comprised within a ∼1 s window, thus establishing an upper range for measurement errors (Fig. 1D, Fig. S1E, E’).

To compare the dynamics of accumulation of Myosin II with RhoA, we then used a RhoA biosensor derived from C-terminus of Anillin fused to GFP to monitor RhoA accumulation (hereon, GFP::AHPH, (Tse et al. 2012)). As previously described (Michaux et al. 2018), we observed a strong spatial and temporal correlation between GFP::AHPH and NMY-2::mKate2, with a median delay of ∼4.5 s of NMY-2::mKate2 with respect to GFP::AHPH (Fig. 1D-E, Movie S2, Fig. S2). These results however only reported on the dynamics of a RhoA biosensor, thus potentially inserting an additional intermediary step in the observed kinetics. To observe directly a sequence of recruitment, we focused on the next step of the cascade. Myosin II activation depends on the phosphorylation level of the Regulatory Light-Chain, which is controlled by phosphorylation by ROCK and dephosphorylation by the Myosin Phosphatase (Karess et al. 1991). We therefore compared the respective dynamics of ROCK and Myosin II, and used a strain coexpressing NMY-2::mKate2 with Rho Kinase fused with GFP at the endogenous genomic locus (GFP::LET-502, (K. R. Bell et al. 2020)). As for GFP::AHPH, we observed a coupled, delayed accumulation of GFP::LET-502 with NMY-2::mKate2. Importantly, we measured a delay of ∼4.5 s between ROCK and Myosin II, not significantly different from the delay between RhoA and Myosin II (Fig. 1D-E, Fig. S1A-D’, Movie S3).

As the inhibition of the Myosin Phosphatase by ROCK has been proposed to contribute to the activation of Myosin II (Piekny and Mains 2002), we also decided to monitor the recruitment dynamics of the MEL-11 regulatory subunit of myosin phosphatase, and generated a MEL-11::GFP CRISPR knockin. In *C. elegans, mel-11* mutants display a range of phenotypes, demonstrating that *mel-11* plays multiple key roles at several stages during embryonic morphogenesis (Piekny and Mains 2002; Wissmann, Ingles, and Mains 1999). As our CRISPR strain displayed no functional phenotype, we concluded that the fusion protein was functional. In early embryos, MEL-11::GFP was cortically enriched, and displayed a dynamic, grainy distribution at the cell surface, accumulating in particular at the cleavage furrow during cell division (Movie S3), as previously proposed based on immunostaining (Piekny and Mains 2002). Co-expressing NMY-2::mKate2 with MEL-11::GFP, we compared their accumulation dynamics. During pulsed contractions at the 2-cell stage, MEL-11 showed dynamic recruitment at the cortex, which did not visually seem to match pulsed contractions (Movie S4). As expected from this observation, attempting to measure the delay between NMY-2::mKate2 and MEL-11::GFP resulted in very poorly synchronized curves (Fig. S1G, Movie S4) and broad dispersion of the delay measurements (Fig. S1G’), further supporting the absence of coordination between MEL-11::GFP cortical recruitment and pulsed contractions. Interestingly, this dispersion differed strongly from the dynamics we observed for all the other players of the activation cascade (Fig. 1D, Fig. S1A-F’), suggesting that Myosin II accumulation is controlled locally by its activation through RhoA and ROCK while the regulation of Myosin II by its phosphatase is not a local process, but instead takes place at the scale of the embryo.

In previous experiments, we used Myosin Heavy-Chain fusion NMY-2::mKate2 as a reporter of Myosin II dynamics. We however wondered if we could distinguish the accumulation dynamics of the Heavy-Chain and Myosin Regulatory Light-Chain, and identify a kinetic signature suggesting a dynamic exchange of the Myosin Regulatory Light-Chain from the Myosin Heavy-Chain. To address this question, we generated a Myosin Regulatory Light-Chain MLC-4 GFP CRISPR knock-in strain. Previous work showed that *mlc-4* is required for proper embryogenesis (Shelton et al. 1999). As the knock-in strain displayed no overt phenotype, we concluded that MLC-4::GFP was functional. In early embryos, MLC-4::GFP displayed a distribution very similar to that of NMY-2::GFP and NMY-2::mKate2. Co-expressing MLC-4::GFP with NMY-2::mKate2, we observed that MLC-4::GFP, and NMY-2::mKate2 were recruited with extremely similar spatial and temporal dynamics, actually reminiscent of our NMY-2::GFP/NMY-2::mKate2 control experiment (Fig. 1D, Fig. S1E-F’, Fig. S2E-F). As previously for NMY-2::GFP and NMY-2::mKate2, we could not measure a statistically significant delay between the recruitment of the two chains (Fig. 1D, Fig. S1F-F’, Movie S4). This observation supports the idea that Myosin II is recruited to the cortex as a hexamer, and that the Myosin Regulatory Light-Chain, once assembled, does not display an exchange dynamics on the Myosin Heavy-Chain, or that this exchange takes place at a timescale outside of our temporal resolution.

We thus dissected the signaling cascade that leads to the recruitment of Myosin to the cortex, precisely defining the delay between Rho, ROCK and Myosin. Importantly, our results looking at Myosin, either with NMY-2::GFP and MLC-4::GFP, show that measurement errors are rather small, and cannot explain the broader range of delays observed with either RhoA/AHPH or ROCK. This variability, be it stochastic or not, shows that contractility shows some level of robustness with respect to the observed range of delays. These observations also raise the question of whether the timing relationships uncovered here are functionally important for pulsed contractility, rather than being passive consequences of molecular turnover.

### Myosin II-ROCK accumulation delay results from Myosin II binding/unbinding kinetics

Taken together, our results demonstrated that the accumulation of Myosin II at the cortex mirrored the accumulation of ROCK, with a time delay of 4.5 s. It remained unclear, however, whether the delay originated from the kinetics of the phosphorylation reaction or if it resulted from the recruitment dynamics of Myosin II. In order to clarify the molecular mechanisms underlying the time delays that we observed in the cascade, we decided to focus on the kinetics of Myosin II accumulation.

We thus performed single-molecule microscopy with particle tracking. We previously established this method to explore kinetics and mobility of fusion proteins expressed at single-molecule levels in the early *C. elegans* embryo (Robin et al. 2014; Michaux et al. 2018). To visualize the dynamics of individual molecules during pulsed contractions, we used an overexpression strain carrying NMY-2 fused with GFP over an endogenous NMY-2 background, and used RNAi against the GFP to specifically decrease the expression of the NMY-2::GFP fusion protein, reverting the initial over-expression to a wild-type phenotype with minute levels of NMY-2::GFP (Fig. 2A-C, Movie S5). Using automated particle tracking, we then tracked individual molecules, and measured the appearance, density and disappearance of molecules in pulsed contractions (Fig. 2F). As in previous studies (Watanabe 2002; Vallotton et al. 2004; Ponti 2004; Robin et al. 2014; Michaux et al. 2018), we assumed that appearance, fraction of disappearing molecules, and density, reported directly on the local cortical binding rate (hereon *k*_*app*_) (Fig. 2D), unbinding rate (*k*_*off*_) (Fig. 2E), and local density, respectively. Finally, we previously showed that, provided that photobleaching rate is low compared to turnover, tracking indeed provides a reasonable estimate of the turnover rate, a condition which we verified for Myosin II (Robin et al. 2014; Michaux et al. 2018). We also accounted for the impact of local cortical shrinkage/expansion, using a routine to follow adaptive regions of interest (Fig. 2C). Briefly, we used the position of molecules to infer local strain and corrected the region of interest so as to proceed to our quantifications in an adaptive frame of reference, thereby removing confounding effects of advection of the cortex (for details, see Michaux *et al*., 2018). These effects, however, only affect the measurements in a very limited manner (<5%).

**Figure 2.**
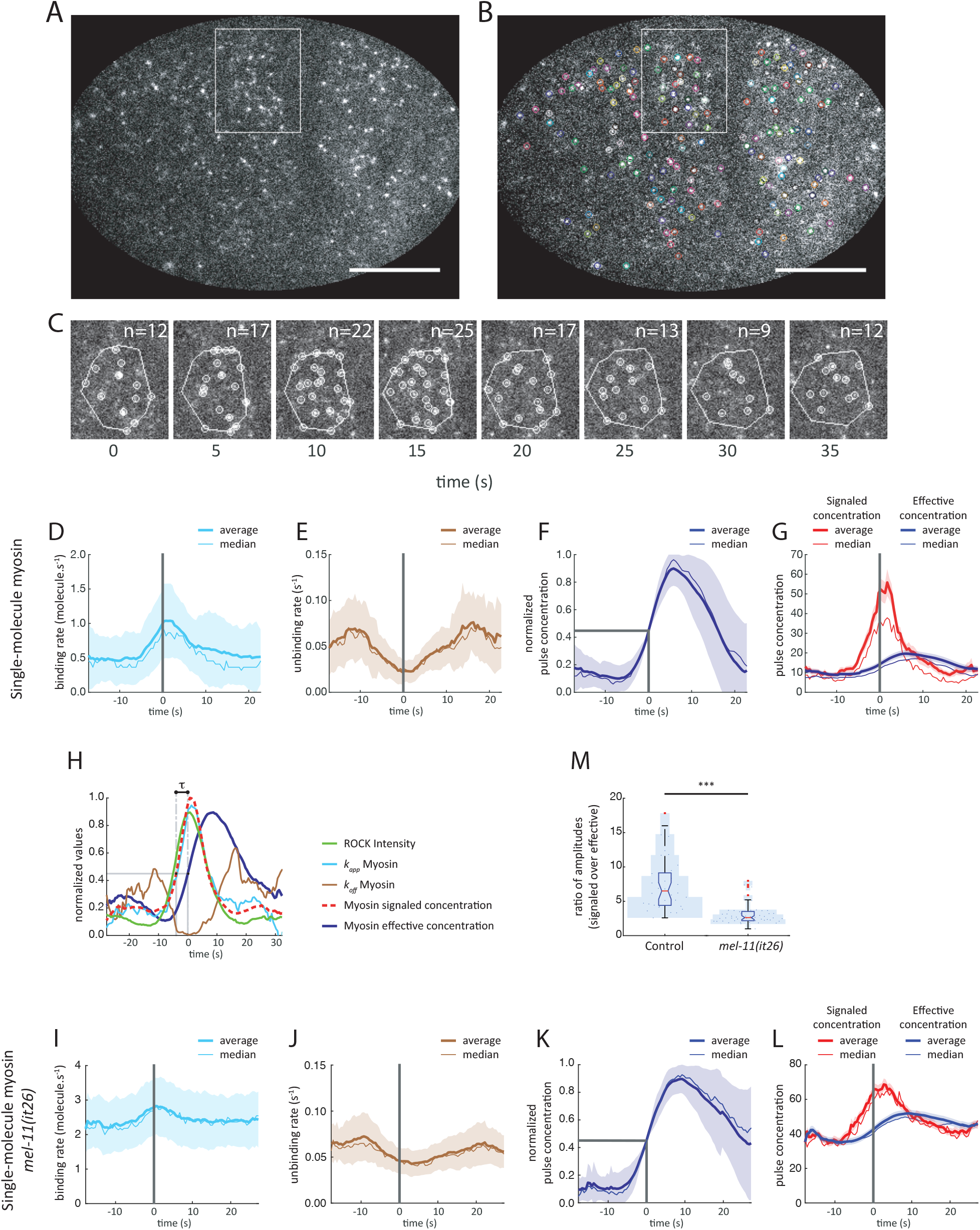
Single-molecule dynamics resolve Myosin binding/unbinding kinetics, revealing the origins of the ROCK/Myosin delay. **(A-B)** Near-TIRF microscopy image of single-molecule of Myosin, scale bar: 10 μm. **(B)** Colored circles: particles detected in image (A) by tracking, each molecule with a circle of a different color. **(C)** Time-lapse of the pulsed contraction in white box in (A). Upper right: number of detected particles. **(D-G)** Measurements from single-molecule microscopy data analysis of Myosin (NMY-2::GFP overexpression). N(embryos) = 5, N(pulses) = 38, thin line: median, thick line: average, shade: (D-F): std, (G): sem **(D)** Binding rate (*K*_*on*_) of Myosin as function of time, **(E)** unbinding rate (*k*_*off*_), **(F)** normalized effective density, **(G)** signaled density (*K*_*on*_/*k*_*off*_, red) and effective density (blue). **(H)** Variation of normalized values over time of ROCK intensity average (Fig. S1B), Myosin intensity average from Supp Fig. 1B, aligned and hence standing for average of Myosin intensity in Fig. 2F (red straight line), Myosin *K*_*on*_ average from Fig. 2D (cyan straight line), Myosin *k*_*off*_ average from Fig. 2E (black straight line), and Myosin signaled density (*K*_*on*_/*k*_*off*_) (magenta dashed line). **(I-L)** Single-molecule tracking of Myosin (NMY-2::GFP overexpression) in Myosin Phosphatase mutant context *mel-11(it26)*. N(embryos) = 9, N(pulses) = 54, thin line: median, thick line: average, (I-K): std, (L): sem **(I)** Binding rate (*K*_*on*_), **(J)** Unbinding rate (K_off_), **(K)** Normalized effective density, **(L)** effective density (blue) and signaled density (*K*_*on*_/*K*_*off*_, red). **(M)** Descriptor to quantify the out-of-equilibrium dynamics of the Myosin, based on the ratio of signaled density amplitude over effective density amplitude for Myosin: (maximum of signaled density – min of signaled density)/(max of effective density – min of effective density) from (G) for the control and (L) for *mel-11(it26)*, ***: p-value ≤ 0.001.

To explore our data, we further hypothesized a very simple kinetic accumulation model, very similar to weakly activated cascades model ((Heinrich, Neel, and Rapoport 2002), see Supplementary Information):

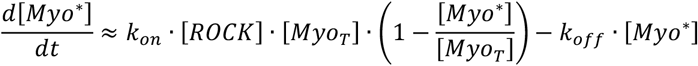

where [*Myo\**] is the concentration of phosphorylated Myosin II, [*Myo*_*T*_] is the total concentration of Myosin II. Under the assumption that the amount of activated Myosin II does not deplete the cytoplasmic stock ([*Myo\**] ≪ [*Myo*_*T*_]), and that ROCK recruitment is not affected by Myosin II, we can write:

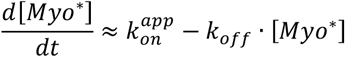

where the effective binding rate (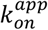, or simply *k*_*app*_) and the unbinding rate (*k*_*off*_) determine the evolution of the cortical concentration of Myosin II (Fig. 3B). Interestingly, under the previous assumptions, *k*_*app*_ is linearly dependent on ROCK concentration.

**Figure 3.**
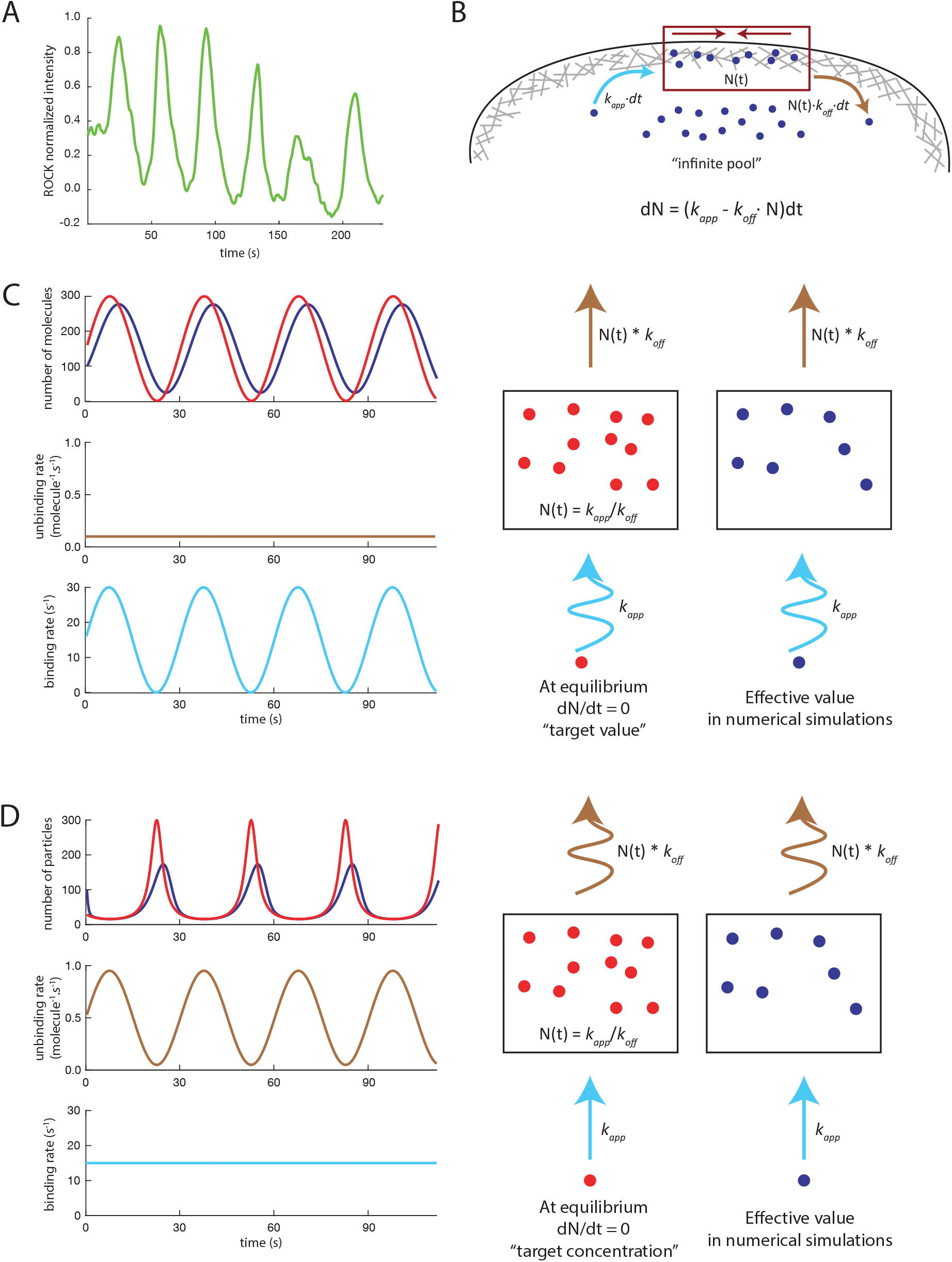
Simple kinetic model supports kinetic control of the delay and highlights the role of unbinding kinetics in shaping Myosin accumulation. **(A)** Normalized intensity variation of ROCK as a function of time, over 6 successive pulsed contractions. **(B)** Schematic of the proposed model for the RhoA activation cascade. **(C)** Blue line: simulation of the effect RhoA periodic variation on the *K*_*on*_ (dark green line), with a constant *k*_*off*_ (light green line), red line: *K*_*on*_/*k*_*off*_. **(D)** Blue line: simulation of the effect RhoA periodic variation on the *k*_*off*_ (light green line), with a constant *K*_*on*_ (dark green line), red line: *K*_*on*_/*k*_*off*_.

Using this equation, and dynamic measurements of effective binding and unbinding rates, we can define an *instantaneous steady-state* concentration [*Myo*^*0*^] for each couple of *k*_*app*_, *k*_*off*_, such that [*Myo*^*0*^] = *k*_*app*_/*k*_*off*_. Biologically, this concentration represents the steady-state concentration that the system would eventually reach if *k*_*app*_ and *k*_*off*_ were to remain constant over time, which we call hereon cortical *signaled concentration*. Importantly, the signaled concentration thus reflects the local signaling intensity of the immediate upstream regulator in the signaling cascade, here the concentration of active ROCK. Fundamentally, comparing the signaled and effective concentrations actually informs on how far out of equilibrium the system lies.

This experimental signaled concentration value therefore integrates modulations of binding and unbinding rate to reflect an activation intensity of the upstream signal in the system. Comparing *signaled concentration* with *effective concentration*, we could therefore measure a delay between the system and its target value, of approximately 4.25 s (Fig. 2H). We then compared the delay between ROCK and Myosin II, measured in the previous section, and the delay from the binding/unbinding kinetics, extracted from our single–molecule measurements. Strikingly, the two results were extremely close, showing that the core of the delay between the accumulation of ROCK and Myosin II essentially (4.25 s out of 4.5) results directly from Myosin II binding/unbinding kinetics (Fig. 2G).

As the signaled concentration should reflect the effect of local ROCK activity, we decided to compare signaled concentration with ROCK/LET-502 local relative intensity. To this end, we aligned the average Myosin II effective concentration of our single-molecule Myosin II dataset with the average relative intensity of NMY-2::mKate2 of our 2-color GFP::LET-502/NMY-2::mKate2 dataset, thus synchronizing the two datasets (Fig. 2H). We then compared the dynamics of the Myosin II signaled concentration with ROCK accumulation. We observed that the two curves are highly overlapping, suggesting that our simple model captures fairly accurately the dynamics of the system.

In essence, our results thus showed that the cortical recruitment of ROCK virtually immediately modulates Myosin II kinetics. In contrast, the evolution of Myosin II concentration towards a dynamically modulated Myosin II signaled concentration, experimentally computed from *k*_*app*_ and *k*_*off*_, takes place with a time constant of several seconds, in a manner *resembling* a capacitor’s charge.

### Exploring the effect of binding/unbinding kinetics using numerical simulations

To better understand how modulations of upstream RhoA/ROCK dynamics affected Myosin II cortical concentration, we turned to numerical simulations. In the cell, RhoA accumulation is sometimes pulsed and unsynchronized, but also sometimes resembles a pseudo-periodic sine wave (Fig. 3A). To get a sense of the effect of the modulation, by RhoA and ROCK, of the binding and unbinding rates, we decided to simulate *k*_*app*_ and *k*_*off*_ as sinusoid modulations (Fig. 3B). Setting binding and unbinding rates, we could then compute both effective and signaled concentrations, and numerically explore how these modulations affected the evolution of the system.

Upon sinusoidal modulations of the binding rate and constant unbinding rate, we observed the emergence of a delay between effective and signaled concentrations (Fig. 3C, blue and red curves, resp.). Unsurprisingly, this delay decreased with increasing *k*_*off*_ (Fig. S3A-C), the unbinding rate acting as a capacitor shifting the phase of the input signal, while the output concentration remained sinusoidal.

An analytical solution of this problem (Supplementary Note 1), interestingly showcases the high-pass filter effect of myosin turnover over upstream pulsed signals, with an attenuation factor 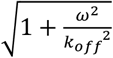.

Conversely, upon constant binding rate and sinusoidal modulations of the unbinding rate, we observed that both effective and signaled concentration displayed a sharp periodic peak around the time *k*_*off*_ was minimal, and were essentially flat and overlapping as *k*_*off*_ was high (Fig. 3D). Interestingly the shape of the curve observed *in vivo* closely resembled this second situation, with *k*_*app*_ constant, varying *k*_*off*_ (Fig. 2G and 3D). Comparing the numerical simulations of the model to the single-molecule results thus suggested that unbinding rate modulation actually played a significant role in shaping Myosin II accumulation.

### Perturbing Myosin II phosphorylation dynamics affects the dynamic range of the system

To test this model, we decided to perturb the unbinding dynamics of Myosin II using *mel-11(it26)*, a temperature-sensitive allele that behaves as a null allele at the restrictive temperature that displays a hypercontractile phenotype during zygote polarization and early cell divisions (Piekny and Mains, 2002). We crossed *mel-11(it26)* with a strain expressing NMY-2::GFP, imaged 2-cell stage embryos at single-molecule levels as described in the previous section, following the quantitative analysis as before (Fig. 2I–L). As expected from its biochemical activity, *mel-11(it26)* displays an increased accumulation of cortical Myosin II (Fig. 2L). We then observed that the average *k*_*app*_ increased ∼3-fold compared to control, while *k*_*off*_ strikingly remained in the same range. Focusing on the dynamics of *k*_*app*_ and *k*_*off*_, we noted that the relative amplitude of the variations in time of both variables, while still present, were much milder in *mel-11(it26)* compared to control embryos (Fig. 2I–F, J–K). Strikingly, the maximum-to-minimum ratio of the signaled concentration was reduced in the mutant context: in the control situation, the signaled concentration increased ∼6-to 7-fold from minimum to maximum signaled concentration, while in contrast, in the mutant, the minimum to maximum signaled concentration displayed only 2-to 3-fold variation (Fig. 2G, L, red curves, Fig. 2M). Similarly, the maximum-to-minimum ratio of the effective concentration was much lower in the *mel-11(it26)* mutant background (∼1.25-fold), compared to control (∼2-fold) (Fig. 2G, L, blue curves). Our results show that *mel-11(it26)*, while displaying a stronger cortical contractility (Piekny and Mains 2002) and higher cortical Myosin II density, also display weaker variations in cortical Myosin II density.

Taken together, these results show that Myosin II dephosphorylation by the Myosin Phosphatase MEL-11 promotes the emergence of strong, well defined pulses. The absence of Myosin Phosphatase thus shifts Myosin II activation out of the dynamic range of the cascade into a saturated regime where Myosin II remains constantly activated. Interestingly however, perturbing Myosin Phosphatase activity did not significantly affect the measured delay between signaled and effective concentration (Fig. 2L), suggesting a degree of a robustness of this delay to perturbations of the phosphatase activity.

### Actin accumulation reflects modulations of polymerization/depolymerization dynamics, and precedes myosin accumulation

During pulsed contractions, in parallel with Myosin II, RhoA regulates F-actin dynamics by directly regulating the formin CYK-1/mDia, an efficient actin filament elongation factor, locally increasing Actin assembly rates (Swan et al. 1998; Costache et al. 2022; Naganathan et al. 2018). In order to explore how the same activation input was transduced across two branches of a cascade, we thus decided to finely describe the temporal dynamics of the F-actin branch of the RhoA cascade.

We first focused on the formin CYK-1. We therefore generated a strain co-expressing CYK-1 fused with GFP at the endogenous genomic locus (CYK-1::GFP, (Costache et al. 2022)), with NMY-2::mKate2, using NMY-2::mKate2 to synchronize our pulses. Using our strategy to quantify cascade kinetics, we measured a delay of ∼4.5 s (Fig. 4A-B, Movie S6). Interestingly, this delay was not significantly different from the RhoA/Myosin II and ROCK/Myosin II delays (Fig. 4A, Fig. S1A–B’, D, D’).

**Figure 4.**
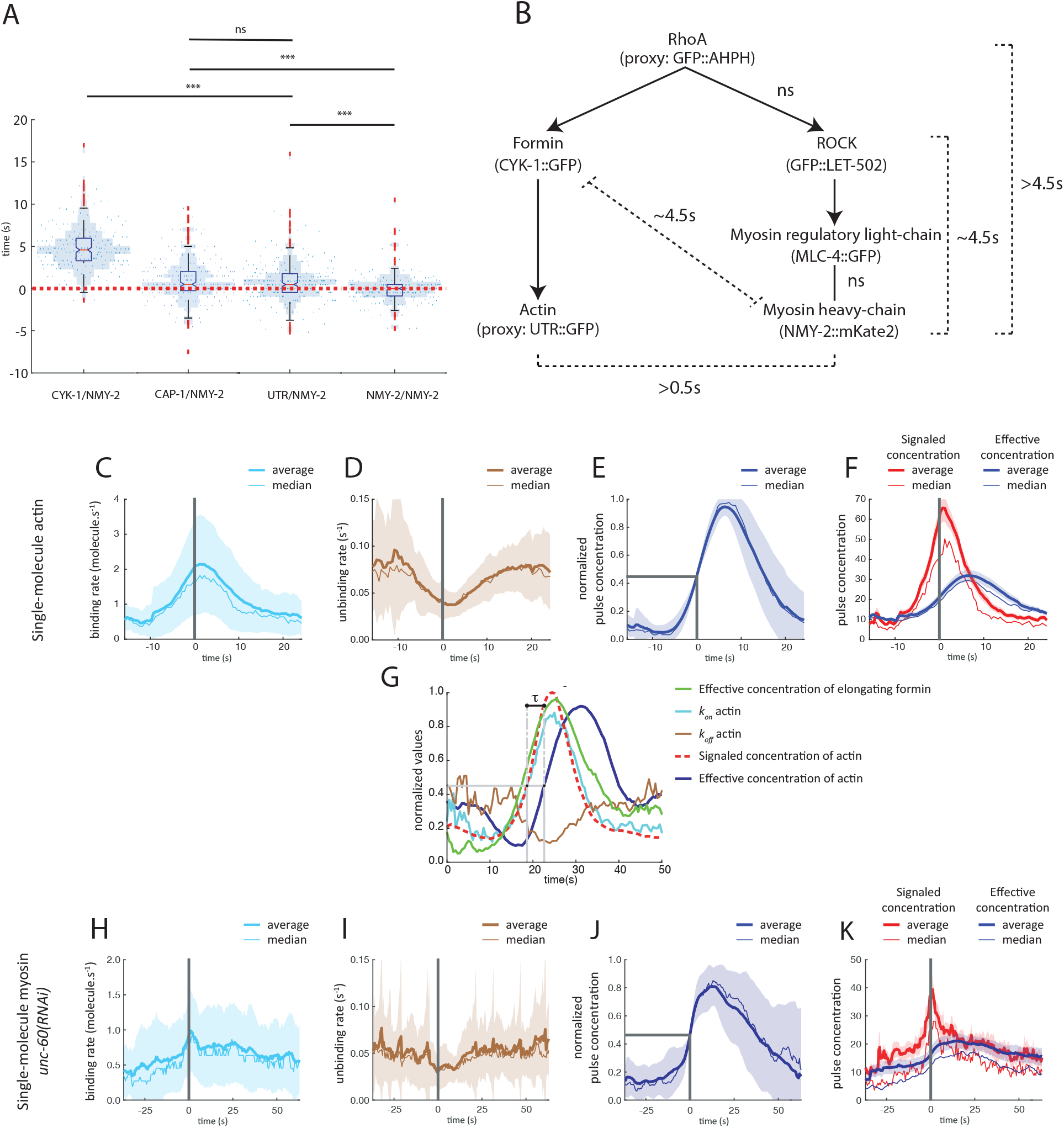
Perturbing Actin dynamics does not alter Myosin *unbinding kinetics*. **(A)** Quantifications of time delay. Blue dots: time delay distribution red over green channel, red dotted line: time delay of 0 s, blue shade: histogram of distribution. N(embryos) = 10, N(pulses) ≥ 200 (See Supplementary Table S1 and S2 for statistical details), ns: not significant, ***: p-value ≤ 0.001. **(B)** Summary of the observed times delay within the RhoA activation cascade. **(C-F)** Single-molecule tracking of Myosin::GFP in cofilin knockdown *unc-60(RNAi)*. N(embryos) = 8, N(pulses) = 39. **(C)** Binding rate (*K*_*on*_) of Myosin over time, **(D)** unbinding rate (*k*_*off*_), **(E)** normalized density, **(F)** effective density (blue) and signaled density (*K*_*on*_/*k*_*off*_, red). **(G-J)** Single-molecule tracking of Actin::GFP. N(embryos) = 6, N(pulses) = 55. (C-J) thin line: median, thick line: mean. (C-E, J-K) shade: std, (F, J) shade: sem. **(G)** Polymerization rate (*K*_*on*_) of Actin over time, **(H)** unbinding rate (*k*_*off*_), **(I)** normalized density, **(J)** effective density (blue) and signaled density (*K*_*on*_/*k*_*off*_, red). **(K)** Variation of normalized intensities over time of Actin average from Supp Fig. 1C (green straight line), Formin super-diffusive population, aligned (red straight line), Actin *K*_*on*_ average from Fig. 4G (cyan straight line), Actin *k*_*off*_ average from Fig. 4H (black straight line), calculated signaled intensity value (*K*_*on*_/*k*_*off*_) (magenta dashed line).

At this point, we suspected that F-actin and Myosin II were synchronously recruited at the cortex. To test this hypothesis, we used a strain co-expressing an Actin reporter –the F-actin-binding *Calponin Homology* domain of Utrophin, fused with GFP (hereon UTR::GFP)– and Myosin II. Surprisingly, we measured a statistically significant delay of ∼0.5 s between F-actin and Myosin II, F-actin appearing briefly before Myosin II (Fig. 4A-B, Fig. S1C, C’, Movie S7). Additionally, as we were not directly observing F-actin, and instead using UTR::GFP to indirectly monitor F-actin accumulation, our results further pointed to the accumulation of F-actin predating the accumulation of Myosin II, with a delay strictly greater than 0.5 s.

To understand the kinetics underlying F-actin accumulation, we wanted to access Actin assembly and depolymerization rates. To this end, we used a strain expressing Actin fused with GFP at single-molecule levels (hereon Actin::GFP), and measured the density of F-actin during pulsed contractions, F-actin binding rate (assembly) and unbinding rate (depolymerization), and F-actin signaled concentration (Fig. 4C–F, (Michaux et al. 2018)). Interestingly, we observed that the ratio between the maximum and minimum binding rates during a pulse was much larger for F-actin (∼5-fold increase, from ∼0.5 to ∼2.5, Fig. 4C) compared to Myosin II (∼2-fold increase, Fig. 2D). In contrast, the comparative variations in *k*_*off*_ displayed the same order of magnitude (Fig. 4D, 2E). This observation suggested that F-actin and Myosin II cortical densities are modulated in very different ways, with a regulation relying largely on assembly for F-actin, while it relies largely on disassembly for Myosin II.

In previous work (Costache et al. 2022), we showed that during pulsed contractions, two populations of formins are recruited at the cortex: *recruited formins*, functionally inactive and immobile, and *elongating formins*, which actively elongate F-actin and move ballistically in the cortex. We then decided to compare single-molecule actin dynamics with the accumulation of the population of elongating formins. To synchronize these F-actin kinetics data with our data on pulse dynamics, we aligned Actin::GFP cortical density with the average curve of our UTR::GFP/NMY-2::mKate2 movies. Similarly, we generated single-molecule microscopy movies of formin fused with GFP (hereon CYK-1::GFP), and used our kinetics data from our CYK-1::GFP/NMY-2::mKate2 data to synchronize our single-molecule dataset with our 2-color dataset. Using these synchronized datasets, we then compared the dynamics of the F-actin signaled concentration with the dynamics of the population of elongating formin (Fig. 4G). As for Myosin II, we observed that the two curves readily overlapped, suggesting that we had captured the essential features of F-actin accumulation.

We finally wondered if F-actin dynamics could be affected by local capping dynamics mediated by the *C. elegans* capping protein CAP-1, which had previously been implicated in pulse dynamics (Naganathan et al. 2018). Using a strain expressing a translational fusion between CAP-1 and GFP at the endogenous genomic locus (hereon CAP-1::GFP), we observed that CAP-1 was cortically enriched and displayed pulsed accumulations among other features. In a strain co-expressing CAP-1::GFP with NMY-2::mKate2, we observed a delay of ∼0.5 s (Fig. 4A, Fig. S1H-H’, Movie S8)between CAP-1 and Myosin II, similar to the delay we had measured between Myosin II and F-actin. This observation suggests that F-actin is dynamically capped at the cortex after F-actin barbed-end are released by formins.

### Slowing down F-actin dynamics affects the dynamic range of the system

To test the effect of actin turnover on pulse dynamics, we decided to perturb and try to predict the impact on the kinetics in the cascade. To slow down actin turnover, we used RNAi against the actin severing protein gene, cofilin *unc-60*. In order to clarify the effects of Cofilin, we first performed *unc-60(RNAi)* in embryos expressing very low levels of a translational fusion of actin with GFP, thus monitoring actin dynamics directly (Robin et al. 2014; Michaux et al. 2018). Using single-molecule measurements of Actin::GFP, we first showed that Cofilin knock-down slowed F-actin turnover (Fig. S4E). We then measured the average pulse period, and observed a shift from a distribution of pulse periods tightly-centered around ∼31 s in control embryos to a much broader distribution centered around 42 s in *unc-60* RNAi (Fig. S4F). Altogether, these results showed that *unc-60(RNAi)* embryos displayed a generally slower F-actin turnover and pulse dynamics.

We then combined *unc-60(RNAi)* with NMY-2::GFP single-molecule microscopy, as in our previous experiments, and measured Myosin II pulse density (Fig. 4J,K), binding rate (*k*_*app*_, Fig. 4H), unbinding rate (*k*_*off*_, Fig. 4I) and signaled concentration (*k*_*app*_/*k*_*off*_, Fig. 4K). As expected from the previous experiment (Fig. S4F), we observed broader pulses and an increased pulse period in *unc-60(RNAi)* compared to control (Fig. S4C-D). Interestingly, the unbinding rate only displayed very mild variations compared to the control. As for the Myosin Phosphatase mutant *mel-11(it26)*, we observed that the ratio between the maximum and minimum signaled concentration was reduced (Fig. 2G, 4K), demonstrating a decreased response to the upstream signal in *unc-60(RNAi)*. Strikingly, neither *unc-60(RNAi)* nor *mel-11(it26)* displayed strong changes in the delay we measured between the rises of signaled concentration and effective concentration.

### Cascade activation delay is required for efficient contractility during pulsed contractions

Our results showed that the two pathways downstream of RhoA activation lead to a recruitment of the actin polymerizing enzyme, initiating an F-actin assembly ∼4.5 s before Myosin II accumulation, eventually resulting in F-Actin accumulation predating Myosin II accumulation by >0.5 s. We therefore wanted to know if this these experimentally observed difference of timing was functionally important for pulsed contractility. However, exploring this question experimentally was difficult, as perturbations would affect multiple aspects of pulsed contractions. In order to more clearly distinguish the effects of the dynamics of the cascade on contractility, we therefore turned to agent-based simulation, to separate the respective effects of timing, duration, and rate of filament elongation.

Using our established computational model of actomyosin networks (Fig. S4A,(Jung, Murrell, and Kim 2015)), we probed the effect on cortical mechanics of the delay between the initiation of formin-induced F-actin elongation in cortex and the recruitment of Myosin II. Using a cortex-like F-actin meshwork (20 μm × 20 μm × 100 nm), we simulated RhoA-driven pulsed contraction by locally modulating the kinetics of Myosin II and F-actin elongation rates, based on previous experimental measurements (Costache et al. 2022). Specifically, to reproduce formin activation, we increased the elongation rate of a fraction of the barbed ends in the RhoA-activated region to 1.2 μm/s, and keeping the barbed ends in this state for ∼10 s, thus generating rapidly elongating Actin filaments and reaching ∼12 μm in length. In parallel, to reproduce myosin activation, we locally turned on Myosin II activity in the RhoA-activated region for 15 s.

To test the impact of delayed Myosin II recruitment on network architecture and the deployment of Myosin II generated forces, we then tuned the duration of the delay between formin activation and Myosin II activation as a variable (Jung, Murrell, and Kim 2015). We observed that simultaneous activation of formins and Myosin II lead to high contraction of both F-actin and Myosin II, and to the formation of long-lasting F-actin aggregates. In contrast, delay at 5 s and 10 s lead to less contraction of the network, avoiding network collapse. To better quantify the mechanical impact of the delay, we measured the network contraction for F-actin and Myosin II (Fig. 5B, C), as well as the sum of the forces and the average force generated locally on the network by the pulsed contraction (Fig. 5C, D). Using these metrics, we observed that Myosin II and F-actin contraction indeed decreased with increasing delay (Fig. 5B, C). Interestingly, we also observed that the force deployed by the contraction evolved non-monotonously and seemed higher when the delay was set at 5 s (Fig. 5D, E).

**Figure 5.**
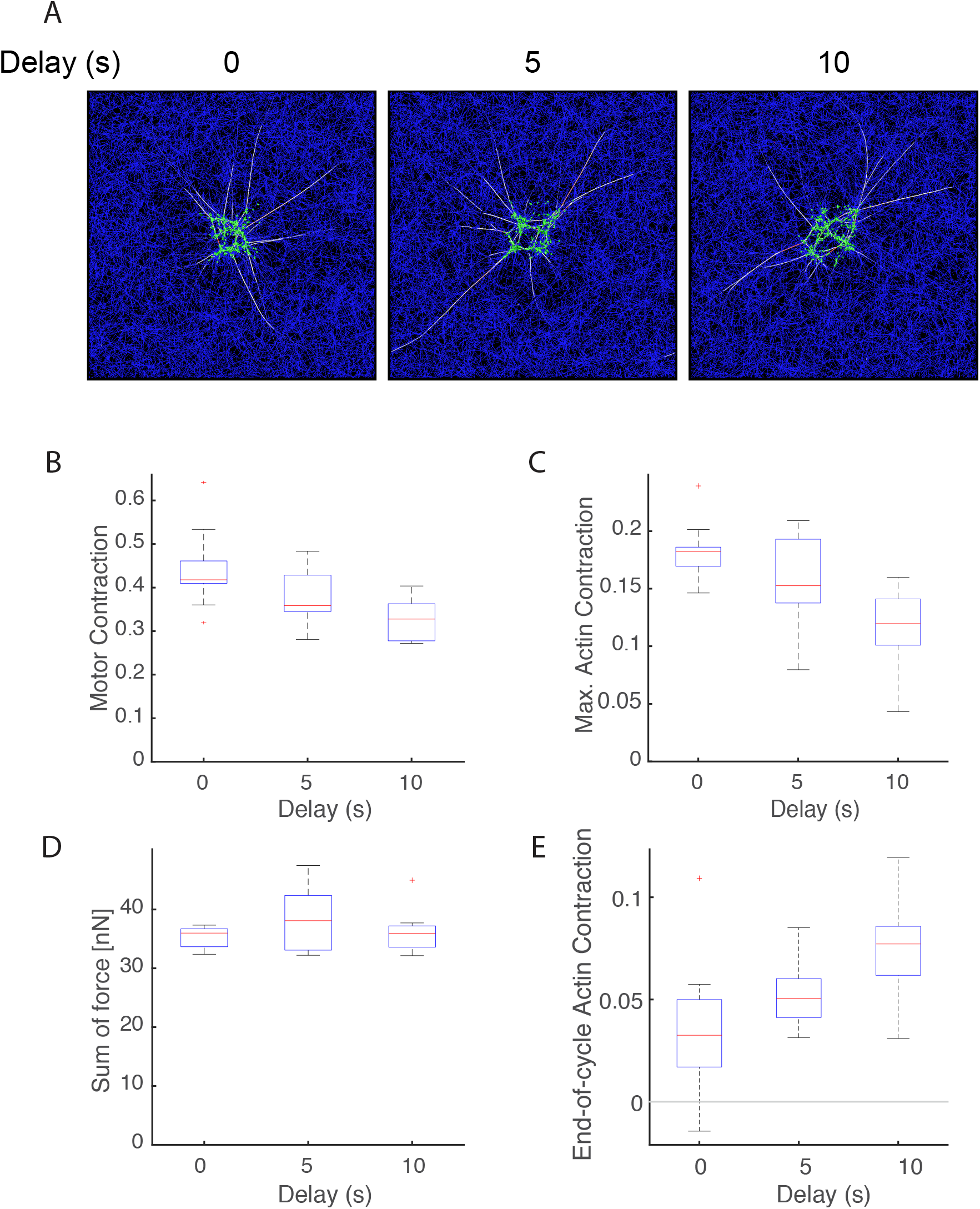
Numerical simulations of actomyosin mechanics reveal that delay between Actin assembly and Myosin accumulation affects force deployment during pulsed contractions. **(A)** Snapshots taken at the end of first myosin activation cycle with an increasing delay between formin-mediated Actin filament elongation and Myosin accumulation. **(B)** Quantification of the extent of motor contraction during pulsed contraction. **(C)** Quantification of actin contraction during pulsed contraction. **(D)** Sum of tensile forces acting and on formin-elongated filaments. **(E)** Average force generated locally and acting on the overall network.

The case without delay showed larger maximum F-actin contraction presumably because there was shorter time for elongated filaments to form cross-linking points before Myosin II motors actively contract Actin filaments. Consistently, the sum of force acting on the network was minimal in the case without the delay (Fig. 5D, E). From these observations, we concluded that the relative timing of Myosin II accumulation and F-actin assembly is actually linked to the contraction and force deployment in the network.

## Discussion

The RhoA pathway driving actomyosin contractility is an evolutionary conserved signaling cascade that drives a broad range of morphogenetic processes, from cytokinesis to cell-cell rearrangements and apical cell constriction to smooth muscle contraction. Depending on the cellular context and RhoA activation modalities, how this signaling cascade is patterned and unfolds can be highly variable, ranging from highly localized, pulsed dynamics to global steady-state activation. Here, coupling live microscopy with image analysis, we identify the enzymatic and kinetic bases that underlie, *in vivo*, the temporal dynamics of the RhoA signaling cascade leading to actomyosin contraction.

Using the RhoA activation cascade in the early *C. elegans* embryo, we followed the temporal dynamics of the cortical accumulation of successive components of cascade. Interestingly, formin, ROCK and RhoA essentially display identical dynamics and delays, suggesting RhoA effectors/binding partners accumulate with similar kinetics. This suggests that our RhoA proxy –like Myosin II– is subject to a binding/unbinding delay and likely lags behind RhoA, and further exploration of the kinetics of AHPH using single-molecule microscopy and particle tracking –as we did for Myosin II and F-actin– should help resolve the actual accumulation kinetics of active RhoA.

Surprisingly, while both RhoA effectors CYK-1 and ROCK/LET-502 accumulate with similar dynamics, F-actin and Myosin II did not appear synchronously at the location of pulsed contractions, F-actin preceding Myosin II by a small yet statistically significant ∼0.5 s delay. Physiologically, this can result from the fact that Myosin II cortical recruitment is dependent on the presence of Actin filaments as binding substrate. In our quantitative analysis, this aspect of Myosin I′I dynamics was implicitly included in the measurement of *k*_*app*_. While this delay is statistically significant on average, individual pulses often display an inversion of recruitment timing. If indeed these inversions reflect genuine biological variability, they suggest that the mechanical output of pulsed contractions is robust to modest variations in the timing of Actin and Myosin recruitment.

Using our single-molecule data analysis, we established a simple model to extract a dynamic *signaled concentration* value –in essence the instantaneous signaling activity of the system– from time-resolved measurements of *k*_*app*_ and *k*_*off*_. Comparing this signaled concentration with effective concentration lets us explore several interesting key properties of the system, and in particular delays, signaling intensity, and out-of-equilibrium dynamics. Importantly, this analysis showed that a single upstream RhoA pulse is decoded into two distinct temporal signals: a rapid polymerization-based response in the actin network, and a delayed, integrative contractile response mediated by myosin turnover.

Importantly, this kinetic model is not specific to the RhoA pathway, and could translate to any number of signaling cascade, as for example MAP kinase cascades operating in the weakly-activated regime (Heinrich, Neel, and Rapoport 2002; Beguerisse-Díaz, Desikan, and Barahona 2016).

Specifically, the model provided a good theoretical framework to understand the molecular bases for the delays we observed along the steps of the cascade. In particular, we could show that Myosin II cortical accumulation relies essentially on the equilibration dynamics under constantly changing *k*_*app*_ and *k*_*off*_. Numerical simulations of RhoA periodic variations and its impact on *k*_*app*_ and *k*_*off*_, independently, showed that *k*_*off*_ modulations under a constant *k*_*app*_ resulted in dynamics extremely closer to the one we observed (Fig. S3).

In order to challenge this model, we experimentally perturbed Actin turnover rate, by depleting the severing factor Cofilin with *unc-60(RNAi)*. While, as expected, this caused an increase in the pulsed contraction period, and abolished *k*_*off*_ variation, it did not affect the kinetic delay for Myosin II inferred from single-molecule measurements. This suggests that Myosin II unbinding is effectively unaffected by Actin turnover. In this context, it therefore seems that Myosin II unbinding operates independently from Actin filament severing.

To further challenge our model, we experimentally perturbed Myosin II dephosphorylation by depleting the Myosin Phosphatase with *mel-11(RNAi)*. Surprisingly, this did not cause a decrease of the average Myosin II unbinding rate. We interpret this result by the fact that Myosin II actually is a low/intermediate duty ratio motor that works cooperatively as mini-filaments. Therefore, the Myosin II unbinding rate might simply represent the unbinding rate of fully phosphorylated Myosin II. Under this assumption, we would expect that an decreased phosphatase activity translates superficially in an increased “available pool” of almost fully phosphorylated Myosin II that could be recruited at the cortex by a weaker phosphorylation activity, thus exclusively affecting the magnitude of the binding rate. This is consistent with the fact that *let-502(sb106RNAi); mel-11(it26)* double mutants are viable: the hypomorph (non-fully functional) *let-502* mutants would in this context work on a larger pool of almost fully phosphorylated myosin, thus resulting in a balance between the two phenotypes. Furthermore, this is consistent with a global, and slow, Myosin Phosphatase activity, which matches our experimental observations regarding MEL-11::GFP spatial localization.

Interestingly, the model also gave indications regarding the molecular bases underlying the magnitude of the response in this dynamic system. In particular, we observed that the RhoA cascade drove myosin concentration in regimes lying far out of equilibrium, with a signaled concentration tuned very far from the effective concentration. In contrast, in Myosin Phosphatase *mel-11(RNAi)* knockdown conditions, the dynamics of the system was much more stable biochemically, suggesting that the signaling cascade likely operated outside of the dynamic range of downstream myosin. The response time of the actomyosin system thus acts as a filter in the time domain on the period of the upstream activation signal: slowly evolving signals keep the system unperturbed, close to the steady state, while fast modulations of the input signal cause the system to oscillate, wandering far out of equilibrium. This behavior is reminiscent of a high-pass filter common in optics, electronics or mechanics, and clearly visible in the analytical resolution of our simplified model representing the system as a forced oscillator under sinusoidal driving signal.

Finally, our agent-based simulations shed some light on how the synchronization of F-actin and Myosin II could actually interplay and subsequently affect Myosin II contractility. We could show that the delay between F-actin and Myosin II may play a physiologically relevant role by promoting long range actomyosin contractility. In this specific context, agent-based simulations of cortical mechanics thus offer an attractive alternative to active gel hydrodynamic models, as they are well suited to capture this type of specifics of emergent properties relying on the detailed architecture of the network.

This work thus reveals an unexpected aspect of signaling cascades, at the crossroads of signaling dynamics, assembly of cytoskeletal architectures, and the mechanical properties of actomyosin networks: kinetic parameters traditionally viewed as passive—binding and unbinding rates—can themselves act as active signal-processing elements to determine both the timing and mechanical outcome of cellular signaling.

## Materials and methods

### Strains

List of strains used in this study can be found in *Supplementary Table 4*. Several strains were provided by the CGC, which is funded by NIH Office of Research Infrastructure Programs (P40 OD010440). MEL-11::GFP transgenic strain was generated by CRISPR as a translational fusion with GFP(65C) at the C-terminus of MEL-11, using a short linker peptide (ACCAGTGGTAGCGGC). CAP-1::GFP transgenic strain was generated by CRISPR as a translational fusion with GFP(65C) at the C-terminus of CAP-1. This strain presents no overt deleterious phenotype, suggesting that the fusion is functional. MLC-4::GFP CRISPR strain is described in Suman, Chauca *et al*. Additional information, including primers and sg are available from the authors upon request.

### *C. elegans* culture, tools and RNAi

Unless specified otherwise, we cultured *C. elegans* at 20°C under standard conditions (Brenner 1974).

Wormbase was used to obtain general information regarding genome, sequences and available tools (Sternberg *et al*, 2024).

*FBR221* hermaphrodites (*mel-11(it26) unc-4(e120) sqt-1(sc13)/mnC1 dpy-10(e128) unc-52(e444)*II; *zuIs45 [nmy-2::NMY-2::GFP + unc-119(+)]* V) were raised at the permissive temperature (15°C). L3-L4 homozygous for *it26* were selected based on phenotype. (*mel-11(it26); zuIs45[nmy-2::gfp])* were placed on Nematode Growth Media (NGM) plates containing 2 mM IPTG for 40 h at 15 °C with HT115 *E. coli* strain expressing GFP RNAi (L4440 GFP RNAi construct), then 1 h prior to the imaging were placed at the restrictive temperature (25 °C).

The RNAi clone targeting *unc-60* was derived from the Ahringer RNAi library (Kamath & Ahringer 2003), and its sequence verified and the plasmid retransformed into HT115(DE3) bacteria. L4 larval stage animals carrying the Myosin II overexpression transgene *zuIs45[nmy-2::gfp]* were placed on Nematode Growth Media (NGM) plates containing 2 mM IPTG for 24 h at 20 °C with HT115 *E. coli* strain expressing GFP RNAi 40 h prior imaging the embryos. Animals were then moved to a plate with 1:1 mix of HT115 *E. coli* strains expressing GFP RNAi and *unc-60* RNAi, and imaged 20 h later.

### Microscopy

Embryos were mounted as described previously (Robin et al., 2014) on 3-well epoxy glass slides (epoxy approx. 20 μm think) under #1.5 coverslips in 2.5 μl of standard Egg Salts containing approx. 100 uniformly sized polystyrene beads (18.7 ± 0.03 μm diameter; no. NT29N; Bangs Laboratories), to achieve uniform compression of the embryos for subsequent TIRF microscopy.

We performed HILO microscopy (Tokunaga, Imamoto, and Sakata-Sogawa 2008) on an inverted Nikon Ti-E microscope, equipped with a motorized TIRF illuminator, a CFI Apo 1.45– NA/100× oil-immersion TIRF objective (Nikon) and a Ti-ND6-PFS-S Perfect Focus unit. Laser illumination at 488 nm from a 50-mW solid-state sapphire laser (Coherent) was delivered by fiber optics to the TIRF illuminator. Images were magnified by 1.5× and collected on an Andor iXon3 897 EMCCD or and Photometrics Prime 95B camera, yielding a pixel size of 107 nm and 73nm, respectively. Microscope and data acquisition parameters were controlled with the NIS software version 4.50.

### Pulsed contractions imaging and image analysis procedures

We imaged at 30% of 90 mW for 488 nm and 561 nm, with 50 ms exposure and no delay between frames, corresponding to an effective 100 ms between two consecutive frames of the same channel. Laser angle was set at 65°. Room temperature was set between 19 and 20,5°C. After acquisition, we used ImageJ to average five consecutive frames, in order to achieve 2 frames per second.

Individual pulses were readily identified in the red channel by the local accumulation of NMY-2. We used ImageJ software (NIH Image, Bethesda, MD) to extract sub-regions containing a single pulsed contraction, as described previously (Michaux et al. 2018; Costache et al. 2022). Individual movies were subsequently loaded and analyzed in Matlab version R2018a. The average image intensity was measured at every time frame for each pulse; the intensity was then normalized:

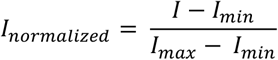

where *I* = mean (Image intensity), *I*_*max*_ = maximal intensity, and *I*_*min*_ = minimal intensity.

Maximum intensity of the red channel is considered as 1 and the immediate precedent minimum is as 0, minimum taken in the 50 preceding frames. Data for the Myosin II intensity were aligned at 0.45, where the slope is maximal and provides the most accurate measure of pulse alignment to fix the time at 0 s for all the red curves. This information is propagated to data from the other channel (green) (which is also normalized to get an intensity between 0 and 1). The average for each distribution (red and green) is plotted with its confidence interval (95%).

For measuring pulse size, we selected manually 5 pulses per embryo and used ImageJ to measure the intensity profile along a line across each pulse for both green and red channels. Comparing pulse sizes, we collected the full width at half maximum (fwhm) and measured the difference between fwhm in green and red.

### Statistical analysis

We performed one-sample Student tests (t-tests) to measure the significance of each delay between paired red and green curves. We used paired-sample Student tests on delays from each channel to measure the significance of the difference between the two delays. *** means p<0.001, **: p<0.01, *: p<0.05, ns: non-significant.

### Single-molecule imaging and Myosin II turnover analysis

We performed single-molecule imaging as described previously (Robin *et al*. 2014). We used RNAi against GFP to reduce the number of imaged fluorescent molecules. We imaged single molecules using 5 % of 90 mW of 488 nm laser, with 500 ms exposure, and no delay between frames. Laser angle was set to 65 °. Room temperature was maintained between 19 and 20,5 °C.

We used Matlab implementation of the Crocker-Grier algorithm by the Kilfoil lab for single-particle tracking (Crocker and Grier 1996; Pelletier et al. 2009). We used the following parameters for all experiments at single-molecule level: particle size, 3 pixels, maximal displacement of the particle between two consecutive frames, 4 pixels and memory to link trajectories in non-consecutive frames, 3 frames. This last parameter allowed the tracking software to look for the particle over multiple frames ensuring robust particle tracking.

Subsequent image analysis was performed in Matlab. Location and time frame of each pulse was loaded in Matlab. At the “seed time”, we defined a Region Of Interest (ROI) corresponding to a convex polygon bounded by molecules detected in the ellipse previously defined in ImageJ (convex envelope of all the particles present in the ROI at the seed time frame). We then displaced this ROI according to the motions of the molecules at the vertices –if present– or by extracting a local velocity field from the motion of the surrounding molecules in a radius of 30 px if the “edge molecule” had disappeared. The ROI thus faithfully described dynamics of the cortex during a pulsed contraction excluding effects from advection caused by the local contraction/expansion of the cortex. We then counted the number of molecules in each frame, the number of molecules appearances (binding rate, *k*_*on*_) and the fraction of molecules disappearing between two consecutive frames (unbinding rate, *k*_*off*_).

Data was normalized to the minimum/maximum number of detected NMY-2 particles, then aligned to 0.45, where the slope is maximal and provides the most accurate measure for pulse alignment. The time of alignment is defined as time = 0 s and is set from the curve of number of molecules and propagated to the curves of binding and unbinding rate. To avoid effects of the beginning and the end of the movie on the binding and unbinding rate, data of number of molecules, for each pulse has been truncated of its first and final three values (replaced by NaN in Matlab). Average for each measurement was plotted with its confidence interval (95%). Equilibrium state value is obtained by division of measured binding and unbinding rates.

We performed the turnover analysis as previously described (Robin et al. 2014). Under the assumption of a single population of particles, the number of molecules *N(t)* within a defined region over time is governed by:

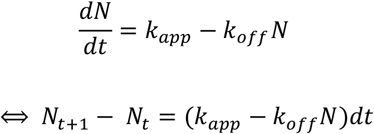

where *k*_*app*_ represents the number of molecules binding (appearing) in a given region per unit time (in molecule/s) and *k*_*off*_ represents the fraction of molecules unbinding per unit time (in s^-1^).

Supposing two different equilibrium states, with distinct *k*_*app*_ and *k*_*off*_ (state i, _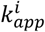_ and _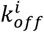_, with equilibrium <u>_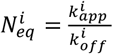_</u>). Then the evolution of the number of molecules between state 1 and state 2, following a step-change from 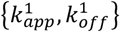to 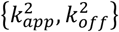 is governed by:

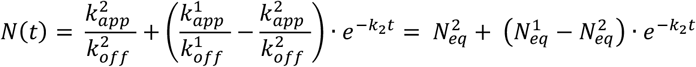

Note that, in our equations, we use concentrations and absolute molecule numbers equivalently, as we account for contractility in tracking our regions of interest. A more generic solution with sinusoidal forcing is proposed in Supplementary Note 1.

To evaluate the difference between signaled concentration and measured density, we computed the fold-difference as follows:

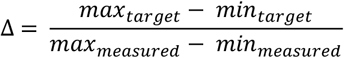

### Comparison between total intensity and dynamic variation at single-molecule level

Normalized average value for ROCK and Myosin II variation in total intensity were extracted from the two-color imaging experiment (GFP::LET-502 and NMY-2::mKate2). The average for Myosin II total intensity was aligned at 0.45 of the maximum with the normalized value of the measured number of molecules in Myosin II single-molecule experiment. The alignment was propagated to the *k*_*on*_, *k*_*off*_ and signaled concentration of the same experiment. On Fig. 2H, we displayed only Myosin II *k*_*on*_, *k*_*off*_ and signaled concentration, and ROCK and Myosin II average normalized intensity.

The normalized average value for formin and Myosin II variation in total intensity were extracted from the two-color imaging experiment (CYK-1::GFP and NMY-2::mKate2). The normalized average value for Myosin II total intensity in this experiment was aligned at 0.45 of the maximum with the normalized average value for Myosin II total intensity of the two-color imaging experiment for F-actin and Myosin II (UTR::GFP and NMY-2::mKate2). The normalized average value for F-actin total intensity was aligned at 0.45 of the maximum with the normalized value of measured number of molecules in F-actin single-molecule experiment. The alignment was propagated to the *k*_*on*_, *k*_*off*_ and the signaled concentration of the same experiment. The normalized average value for formin total intensity in the two-color imaging experiment was aligned at 0.45 of the maximum with the normalized number of superdiffusive formin in formin single-molecule experiment (data published in Fig. 2A-B (Costache et al. 2022). On Fig. 4G, we only displayed the average for F-actin *k*_*on*_ and *k*_*off*_, F-actin signaled concentration, normalized superdiffusive formin population and normalized actin intensity.

### Overview of the computational model of actomyosin mechanics

We used a well-established agent-based model of actomyosin networks based on the Langevin equation (Li et al. 2017; Jung, Murrell, and Kim 2015; T. Kim et al. 2009; Mak et al. 2016). In the model, actin filament (F-actin), motor, and actin crosslinker proteins (ACPs) are coarse-grained using cylindrical segments. The motions of all the cylindrical segments are governed by the Langevin equation for Brownian dynamics. Deterministic forces in the Langevin equation include bending and extensional forces that maintain equilibrium angles formed by segments and the equilibrium lengths of segments, respectively, as well as a repulsive force acting between neighboring pairs of segments for considering volume-exclusion effects.

The formation of F-actin is initiated by a nucleation event, followed by polymerization at the barbed end and depolymerization at the pointed end. ACPs bind to F-actin without preference for cross-linking angles at a constant rate and also unbind from F-actin at a force-dependent rate determined by Bell’s law (G. I. Bell 1978). Each arm of motors binds to F-actin at a constant rate, and it then walks toward the barbed end of F-actin or unbinds from F-actin at force-dependent rates determined by the parallel cluster model (Erdmann, Albert, and Schwarz 2013; Erdmann and Schwarz 2012). For all simulations in this study, we used a thin computational domain (20 × 20 × 0.1 μm) with periodic boundary conditions only in x and y directions. In z direction, the boundaries of the domain exert repulsive forces on elements that moved beyond the boundaries. At the beginning of each simulation, a thin actin network is formed via self-assembly of F-actin and ACP.

For implementing RhoA activation, the domain is divided into 16 subdomains (4×4 in x and y directions). Every 30 s, one of the subdomains is randomly selected and then activated. In the activated subdomain, a fraction of the barbed ends of F-actins are randomly chosen and then undergo faster polymerization by a factor, *ρ*_f_, for the duration of *τ*_f_. With the reference values of *ρ*_f_ = 10 and *τ*_f_ = 10 s, F-actins are elongated by ∼10 μm on average. After the time delay of *d*_M_, motors in the activated subdomain are allowed to self-assemble into thick filament structures for the duration of *τ*_M_. The reference values of *d*_M_ and *τ*_M_ are 5 s and 15 s, respectively. These active motors in the form of thick filaments can contract the part of the network in the activated subdomain. Once they become inactive after *τ*_M_, the motors are disassembled into monomers that cannot bind to F-actin.

## References

Barrett, Kathy, Maria Leptin, and Jeffrey Settleman. 1997. “The Rho GTPase and a Putative RhoGEF Mediate a Signaling Pathway for the Cell Shape Changes in Drosophila Gastrulation.” Cell 91 (7): 905–15. doi:10.1016/s0092-8674(00)80482-1.

Beguerisse-Díaz, Mariano, Radhika Desikan, and Mauricio Barahona. 2016. “Linear Models of Activation Cascades: Analytical Solutions and Coarse-Graining of Delayed Signal Transduction.” Journal of The Royal Society Interface 13 (121): 20160409. doi:10.1098/rsif.2016.0409.

Bell, George I. 1978. “Models for the Specific Adhesion of Cells to Cells.” Science 200 (4342): 618–27. doi:10.1126/science.347575.

Bell, Kathryn Rehain, Michael E. Werner, Anusha Doshi, Daniel B. Cortes, Adam Sattler, Thanh Vuong-Brender, Michel Labouesse, and Amy Shaub Maddox. 2020. “Novel Cytokinetic Ring Components Drive Negative Feedback in Cortical Contractility.” Molecular Biology of the Cell 31 (15): 1623–36. doi:10.1091/mbc.e20-05-0304.

Brenner, S. 1974. “THE GENETICS OF CAENORHABDITIS ELEGANS.” Genetics 77 (1): 71–94. doi:10.1093/genetics/77.1.71.

Cavanaugh, Kate E., Michael F. Staddon, Edwin Munro, Shiladitya Banerjee, and Margaret L. Gardel. 2020. “RhoA Mediates Epithelial Cell Shape Changes via Mechanosensitive Endocytosis.” Developmental Cell 52 (2): 152-166.e5. doi:10.1016/j.devcel.2019.12.002.

Costache, Vlad, Serena Prigent Garcia, Camille N Plancke, Jing Li, Simon Begnaud, Shashi Kumar Suman, Anne-Cécile Reymann, Taeyoon Kim, and Francois B Robin. 2022. “Rapid Assembly of a Polar Network Architecture Drives Efficient Actomyosin Contractility.” Cell Reports 39 (9): 110868. doi:10.1016/j.celrep.2022.110868.

Crocker, J C, and D G Grier. 1996. “Methods of Digital Video Microscopy for Colloidal Studies.” Journal of Colloid and Interface Science 179 (1): 298–310. doi:10.1006/jcis.1996.0217.

de Seze J, Bongaerts M, Boulevard B, Coppey M. 2025. “Optogenetic control of a GEF of RhoA uncovers a signaling switch from retraction to protrusion.” Elife. 12:RP93180. doi: 10.7554/eLife.93180.

Dickinson, Daniel J., Francoise Schwager, Lionel Pintard, Monica Gotta, and Bob Goldstein. 2017. “A Single-Cell Biochemistry Approach Reveals PAR Complex Dynamics during Cell Polarization.” Developmental Cell 42 (4): 416-434.e11. doi:10.1016/j.devcel.2017.07.024.

Diogon, M, F Wissler, S Quintin, Y Nagamatsu, S Sookhareea, F Landmann, H Hutter, N Vitale, and M Labouesse. 2007. “The RhoGAP RGA-2 and LET-502/ROCK Achieve a Balance of Actomyosin-Dependent Forces in C. Elegans Epidermis to Control Morphogenesis.” Development 134 (13): 2469–79. doi:10.1242/dev.005074.

Eaton, S, P Auvinen, L Luo, Y N Jan, and K Simons. 1995. “CDC42 and Rac1 Control Different Actin-Dependent Processes in the Drosophila Wing Disc Epithelium.” The Journal of Cell Biology 131 (1): 151–64. doi:10.1083/jcb.131.1.151.

Erdmann, Thorsten, Philipp J. Albert, and Ulrich S. Schwarz. 2013. “Stochastic Dynamics of Small Ensembles of Non-Processive Molecular Motors: The Parallel Cluster Model.” The Journal of Chemical Physics 139 (17): 175104. doi:10.1063/1.4827497.

Erdmann, Thorsten, and Ulrich S. Schwarz. 2012. “Stochastic Force Generation by Small Ensembles of Myosin II Motors.” Physical Review Letters 108 (18): 188101. doi:10.1103/physrevlett.108.188101.

Ferrell, James E. 2013. “Feedback Loops and Reciprocal Regulation: Recurring Motifs in the Systems Biology of the Cell Cycle.” Current Opinion in Cell Biology 25 (6): 676–86. doi:10.1016/j.ceb.2013.07.007.

Gally, Christelle, Frédéric Wissler, Hala Zahreddine, Sophie Quintin, Frédéric Landmann, and Michel Labouesse. 2009. “Myosin II Regulation during C. Elegans Embryonic Elongation:LET-502/ROCK, MRCK-1 and PAK-1, Three Kinases with Different Roles.” Development 136 (18): 3109–19. doi:10.1242/dev.039412.

Gérard, Claude, and Albert Goldbeter. 2012. “Entrainment of the Mammalian Cell Cycle by the Circadian Clock: Modeling Two Coupled Cellular Rhythms.” PLoS Computational Biology 8 (5): e1002516. doi:10.1371/journal.pcbi.1002516.

Goldbeter, Albert. 1996. Biochemical Oscillations and Cellular Rhythms: The Molecular Bases of Periodic and Chaotic Behavior. Cambridge Univ. Press. Cambridge Univ. Press.

Hariharan, I K, K Q Hu, H Asha, A Quintanilla, R M Ezzell, and J Settleman. 1995. “Characterization of Rho GTPase Family Homologues in Drosophila Melanogaster: Overexpressing Rho1 in Retinal Cells Causes a Late Developmental Defect.” The EMBO Journal 14 (2): 292–302. doi:10.1002/j.1460-2075.1995.tb07003.x.

He, Li, Xiaobo Wang, Ho Lam Tang, and Denise J. Montell. 2010. “Tissue Elongation Requires Oscillating Contractions of a Basal Actomyosin Network.” Nature Cell Biology 12 (12): 1133–42. doi:10.1038/ncb2124.

Heinrich, Reinhart, Benjamin G. Neel, and Tom A. Rapoport. 2002. “Mathematical Models of Protein Kinase Signal Transduction.” Molecular Cell 9 (5): 957–70. doi:10.1016/s1097-2765(02)00528-2.

Huang, C Y, and J E Ferrell. 1996. “Ultrasensitivity in the Mitogen-Activated Protein Kinase Cascade.” Proceedings of the National Academy of Sciences 93 (19): 10078–83. doi:10.1073/pnas.93.19.10078.

Jr., Daniel E. Koshland, Albert Goldbeter, and Jeffry B. Stock. 1982. “Amplification and Adaptation in Regulatory and Sensory Systems.” Science 217 (4556): 220–25. doi:10.1126/science.7089556.

Jung, Wonyeong, Michael P. Murrell, and Taeyoon Kim. 2015. “F-Actin Cross-Linking Enhances the Stability of Force Generation in Disordered Actomyosin Networks.” Computational Particle Mechanics 2 (4): 317–27. doi:10.1007/s40571-015-0052-9.

Kholodenko, Boris N. 2006. “Cell-Signalling Dynamics in Time and Space.” Nature Reviews Molecular Cell Biology 7 (3): 165–76. doi:10.1038/nrm1838.

Kim, H Y, and L A Davidson. 2011. “Punctuated Actin Contractions during Convergent Extension and Their Permissive Regulation by the Non-Canonical Wnt-Signaling Pathway.” Journal of Cell Science 124 (4): 635–46. doi:10.1242/jcs.067579.

Kim, Taeyoon, Wonmuk Hwang, Hyungsuk Lee, and Roger D. Kamm. 2009. “Computational Analysis of Viscoelastic Properties of Crosslinked Actin Networks.” PLoS Computational Biology 5 (7): e1000439. doi:10.1371/journal.pcbi.1000439.

Kovar, David R, and Thomas D Pollard. 2004. “Insertional Assembly of Actin Filament Barbed Ends in Association with Formins Produces Piconewton Forces.” Proceedings of the National Academy of Sciences of the United States of America 101 (41): 14725–30. doi:10.1073/pnas.0405902101.

Li, Jing, Thomas Biel, Pranith Lomada, Qilin Yu, and Taeyoon Kim. 2017. “Buckling-Induced F-Actin Fragmentation Modulates the Contraction of Active Cytoskeletal Networks.” Soft Matter 13 (17): 3213–20. doi:10.1039/c6sm02703b.

Ma, Wenzhe, Ala Trusina, Hana El-Samad, Wendell A. Lim, and Chao Tang. 2009. “Defining Network Topologies That Can Achieve Biochemical Adaptation.” Cell 138 (4): 760–73. doi:10.1016/j.cell.2009.06.013.

Maître, Jean-Léon, Ritsuya Niwayama, Hervé Turlier, François Nédélec, and Takashi Hiiragi. 2015. “Pulsatile Cell-Autonomous Contractility Drives Compaction in the Mouse Embryo.” Nature Cell Biology 17 (7): 849–55.

Mak, Michael, Muhammad H. Zaman, Roger D. Kamm, and Taeyoon Kim. 2016. “Interplay of Active Processes Modulates Tension and Drives Phase Transition in Self-Renewing, Motor-Driven Cytoskeletal Networks.” Nature Communications 7 (1): 10323. doi:10.1038/ncomms10323.

Markevich, Nick I., Jan B. Hoek, and Boris N. Kholodenko. 2004. “Signaling Switches and Bistability Arising from Multisite Phosphorylation in Protein Kinase Cascades.” The Journal of Cell Biology 164 (3): 353–59. doi:10.1083/jcb.200308060.

Martin, Adam C, Matthias Kaschube, and Eric F Wieschaus. 2009. “Pulsed Contractions of an Actin-Myosin Network Drive Apical Constriction.” Nature 457 (7228): 495–99. doi:10.1038/nature07522.

Michaux, Jonathan B., François B. Robin, William M. McFadden, and Edwin M. Munro. 2018. “Excitable RhoA Dynamics Drive Pulsed Contractions in the Early C. Elegans Embryo.” Journal of Cell Biology 217 (12): 4230–52. doi:10.1083/jcb.201806161.

Mori, Yoichiro, Alexandra Jilkine, and Leah Edelstein-Keshet. 2008. “Wave-Pinning and Cell Polarity from a Bistable Reaction-Diffusion System.” Biophysical Journal 94 (9): 3684–97. doi:10.1529/biophysj.107.120824.

Motegi, Fumio, and Asako Sugimoto. 2006. “Sequential Functioning of the ECT-2 RhoGEF, RHO-1 and CDC-42 Establishes Cell Polarity in Caenorhabditis Elegans Embryos.” Nature Cell Biology 8 (9): 978–85. doi:10.1038/ncb1459.

Munjal, Akankshi, Jean-Marc Philippe, Edwin Munro, and Thomas Lecuit. 2015. “A Self-Organized Biomechanical Network Drives Shape Changes during Tissue Morphogenesis.” Nature 524 (7565): 351–55. doi:10.1038/nature14603.

Naganathan, Sundar Ram, Sebastian Fürthauer, Josana Rodriguez, Bruno Thomas Fievet, Frank Jülicher, Julie Ahringer, Carlo Vittorio Cannistraci, and Stephan W Grill. 2018. “Morphogenetic Degeneracies in the Actomyosin Cortex.” ELife 7 (October): e37677. doi:10.7554/elife.37677.

Nance, J. 2003. “C. Elegans PAR-3 and PAR-6 Are Required for Apicobasal Asymmetries Associated with Cell Adhesion and Gastrulation.” Development 130 (22): 5339–50. doi:10.1242/dev.00735.

Novák, Béla, and John J Tyson. 2008. “Design Principles of Biochemical Oscillators.” Nature Reviews Molecular Cell Biology 9 (12): 981–91. doi:10.1038/nrm2530.

Pelletier, Vincent, Naama Gal, Paul Fournier, and Maria Kilfoil. 2009. “Microrheology of Microtubule Solutions and Actin-Microtubule Composite Networks.” Physical Review Letters 102 (18): 188303. doi:10.1103/physrevlett.102.188303.

Piekny, Alisa J, and Paul E Mains. 2002. “Rho-Binding Kinase (LET-502) and Myosin Phosphatase (MEL-11) Regulate Cytokinesis in the Early Caenorhabditis Elegans Embryo.” Journal of Cell Science 115 (Pt 11): 2271–82. http://pubmed.gov/12006612.

Pomerening, Joseph R., Eduardo D. Sontag, and James E. Ferrell. 2003. “Building a Cell Cycle Oscillator: Hysteresis and Bistability in the Activation of Cdc2.” Nature Cell Biology 5 (4): 346–51. doi:10.1038/ncb954.

Ponti, A. 2004. “Two Distinct Actin Networks Drive the Protrusion of Migrating Cells.” Science 305 (5691): 1782–86. doi:10.1126/science.1100533.

Robin François B, William M McFadden, Baixue Yao, and Edwin M Munro. 2014. “Single-Molecule Analysis of Cell Surface Dynamics in Caenorhabditis Elegans Embryos.” Nature Methods 11 (6): 677–82. doi:10.1038/nmeth.2928.

Schonegg, Stephanie, Alexandru T Constantinescu, Carsten Hoege, and Anthony A Hyman. 2007. “The Rho GTPase-Activating Proteins RGA-3 and RGA-4 Are Required to Set the Initial Size of PAR Domains in Caenorhabditis Elegans One-Cell Embryos.” Proceedings of the National Academy of Sciences of the United States of America 104 (38): 14976–81. doi:10.1073/pnas.0706941104.

Solon, Jerome, Aynur Kaya-Copur, Julien Colombelli, and Damian Brunner. 2009. “Pulsed Forces Timed by a Ratchet-like Mechanism Drive Directed Tissue Movement during Dorsal Closure.” CELL 137 (7): 1331–42. doi:10.1016/j.cell.2009.03.050.

Staddon, Michael F., Kate E. Cavanaugh, Edwin M. Munro, Margaret L. Gardel, and Shiladitya Banerjee. 2019. “Mechanosensitive Junction Remodeling Promotes Robust Epithelial Morphogenesis.” Biophysical Journal 117 (9): 1739–50. doi:10.1016/j.bpj.2019.09.027.

Sternberg PW, Van Auken K, Wang Q, Wright A, Yook K, Zarowiecki M, Arnaboldi V, Becerra A, Brown S, Cain S, Chan J, Chen WJ, Cho J, Davis P, Diamantakis S, Dyer S, Grigoriadis D, Grove CA, Harris T, Howe K, Kishore R, Lee R, Longden I, Luypaert M, Müller HM, Nuin P, Quinton-Tulloch M, Raciti D, Schedl T, Schindelman G, Stein L. 2024. “WormBase024: status and transitioning to Alliance infrastructure.” Genetics 227 (1). doi:10.1093/genetics/iyae050.

Swan, K A, A F Severson, J C Carter, P R Martin, H Schnabel, R Schnabel, and B Bowerman. 1998. “Cyk-1: A C. Elegans FH Gene Required for a Late Step in Embryonic Cytokinesis.” Journal of Cell Science 111 (Pt 14) (July): 2017–27. http://pubmed.gov/9645949.

Tan, Pei Yi, and Ronen Zaidel-Bar. 2015. “Transient Membrane Localization of SPV-1 Drives Cyclical Actomyosin Contractions in the C. Elegans Spermatheca.” Current Biology 25 (2): 141–51. doi:10.1016/j.cub.2014.11.033.

Tokunaga, Makio, Naoko Imamoto, and Kumiko Sakata-Sogawa. 2008. “Highly Inclined Thin Illumination Enables Clear Single-Molecule Imaging in Cells.” Nature Methods 5 (2): 159–61. doi:10.1038/nmeth1171.

Tse, Yu Chung, Michael Werner, Katrina M Longhini, Jean-Claude Labbé, Bob Goldstein, and Michael Glotzer. 2012. “RhoA Activation during Polarization and Cytokinesis of the Early Caenorhabditis Elegans Embryo Is Differentially Dependent on NOP-1 and CYK-4.” Molecular Biology of the Cell 23 (20): 4020–31. doi:10.1091/mbc.e12-04-0268.

Valon L, Etoc F, Remorino A, di Pietro F, Morin X, Dahan M, Coppey M. 2015. “Predictive Spatiotemporal Manipulation of Signaling Perturbations Using Optogenetics.” Biophys J. 109(9):1785–97. doi: 10.1016/j.bpj.2015.08.042.

Vallotton, Pascal, Stephanie L Gupton, Clare M Waterman-Storer, and Gaudenz Danuser. 2004. “Simultaneous Mapping of Filamentous Actin Flow and Turnover in Migrating Cells by Quantitative Fluorescent Speckle Microscopy.” Proceedings of the National Academy of Sciences of the United States of America 101 (26): 9660–65. doi:10.1073/pnas.0300552101.

Wallace, Andre G., Hamidah Raduwan, John Carlet, and Martha C. Soto. 2018. “The RhoGAP HUM-7/Myo9 Integrates Signals to Modulate RHO-1/RhoA during Embryonic Morphogenesis in Caenorhabditis Elegans.” Development 145 (23): dev168724. doi:10.1242/dev.168724.

Watanabe, N. 2002. “Single-Molecule Speckle Analysis of Actin Filament Turnover in Lamellipodia.” Science 295 (5557): 1083–86. doi:10.1126/science.1067470.

Wissmann, A, J Ingles, and P E Mains. 1999. “The Caenorhabditis Elegans Mel-11 Myosin Phosphatase Regulatory Subunit Affects Tissue Contraction in the Somatic Gonad and the Embryonic Epidermis and Genetically Interacts with the Rac Signaling Pathway.” Developmental Biology 209 (1): 111–27. doi:10.1006/dbio.1999.9242.

Wissmann, A, J Ingles, J D McGhee, and P E Mains. 1997. “Caenorhabditis Elegans LET-502 Is Related to Rho-Binding Kinases and Human Myotonic Dystrophy Kinase and Interacts Genetically with a Homolog of the Regulatory Subunit of Smooth Muscle Myosin Phosphatase to Affect Cell Shape.” Genes & Development 11 (4): 409–22. doi:10.1101/gad.11.4.409.

Yamaguchi, Hiroto Q., Koji L. Ode, and Hiroki R. Ueda. 2021. “A Design Principle for Posttranslational Chaotic Oscillators.” IScience 24 (1): 101946. doi:10.1016/j.isci.2020.101946.

