## Supplementary Materials for "Kinetic Control of Out-Of-Equilibrium Dynamics in the RhoA Signaling Cascade Shapes Actomyosin Contractility"

The following document includes:

Supplementary Fig. 1
Supplementary Fig. 2
Supplementary Fig. 3
Supplementary Fig. 4
Supplementary Table 1
Supplementary Table 2
Supplementary Table 3
Supplementary Table 4
Supplementary Note 1

### Figure S1

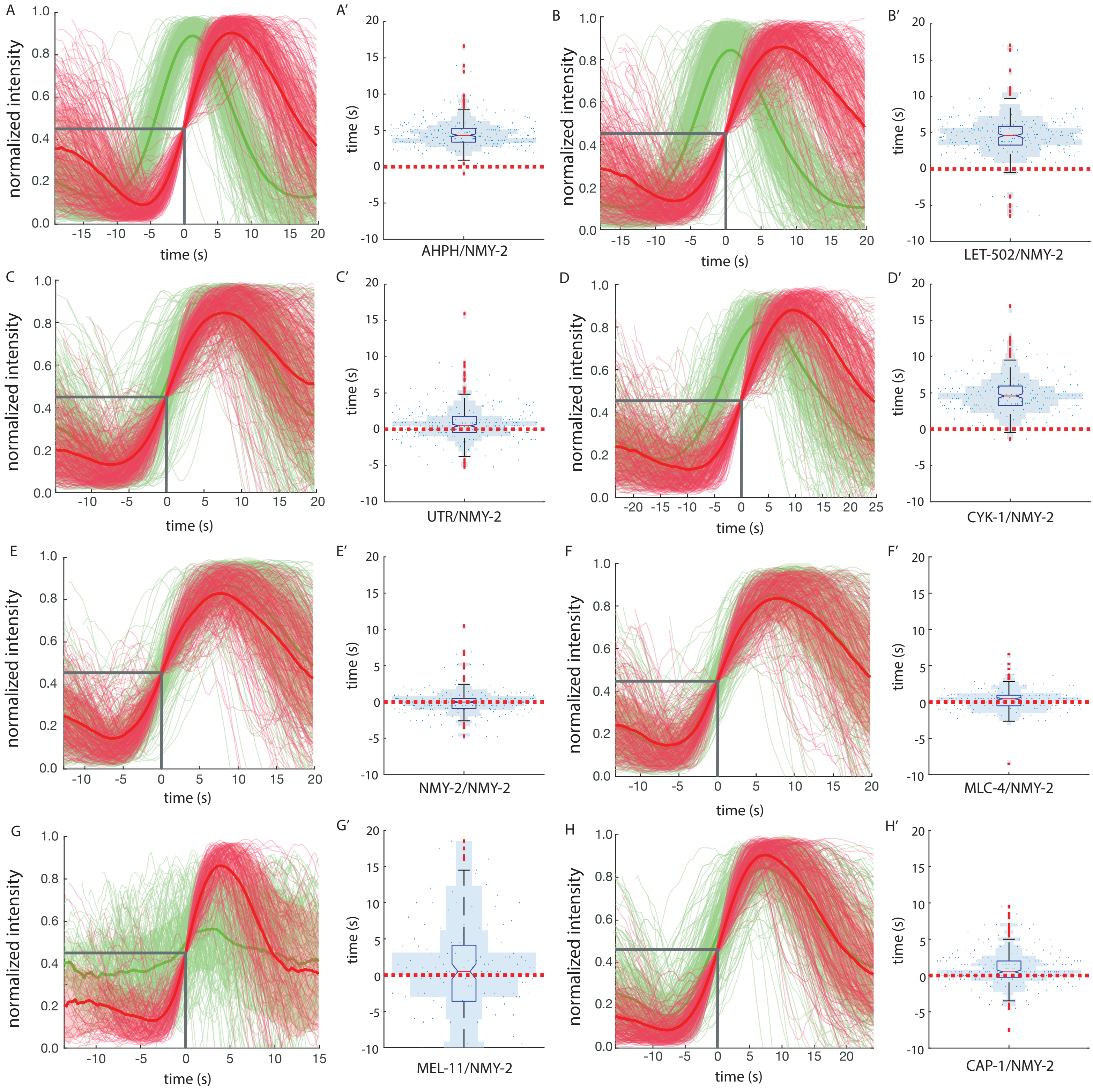

A
A′
B
B′
C
C′
D
D′
E
E′
F
F′
G
G′
H
H′
normalized intensity
time (s)
AHPH/NMY-2
LET-502/NMY-2
UTR/NMY-2
CYK-1/NMY-2
NMY-2/NMY-2
MLC-4/NMY-2
MEL-11/NMY-2
CAP-1/NMY-2

**Supplemental figure 1. Most of the time delay observed between RhoA and Myosin can be explained by the delay between ROCK and Myosin. (A-G')** Time delay between in red Myosin heavy-chain NMY-2 and in green RhoA's proxy AHPH (A-A'), the ROCK LET-502 (B-B'), Actin's proxy UTR (C-C'), the Formin CYK-1 (D-D'), the control with the Myosin overexpression (E-E'), the Myosin regulatory light-chain MLC-4 (F-F'), the Myosin Phosphatase MEL-11 (G-G'), Actin capping-protein CAP-1 (H-H'). **(A-H)** Fainted lines: distribution of normalized intensity through time, solid line: mean, shade: std. **(A'-H')** Blue dots: time delay distribution red over green channel, red dotted line: time delay of 0 s, blue shade: histogram of distribution.

### Figure S2

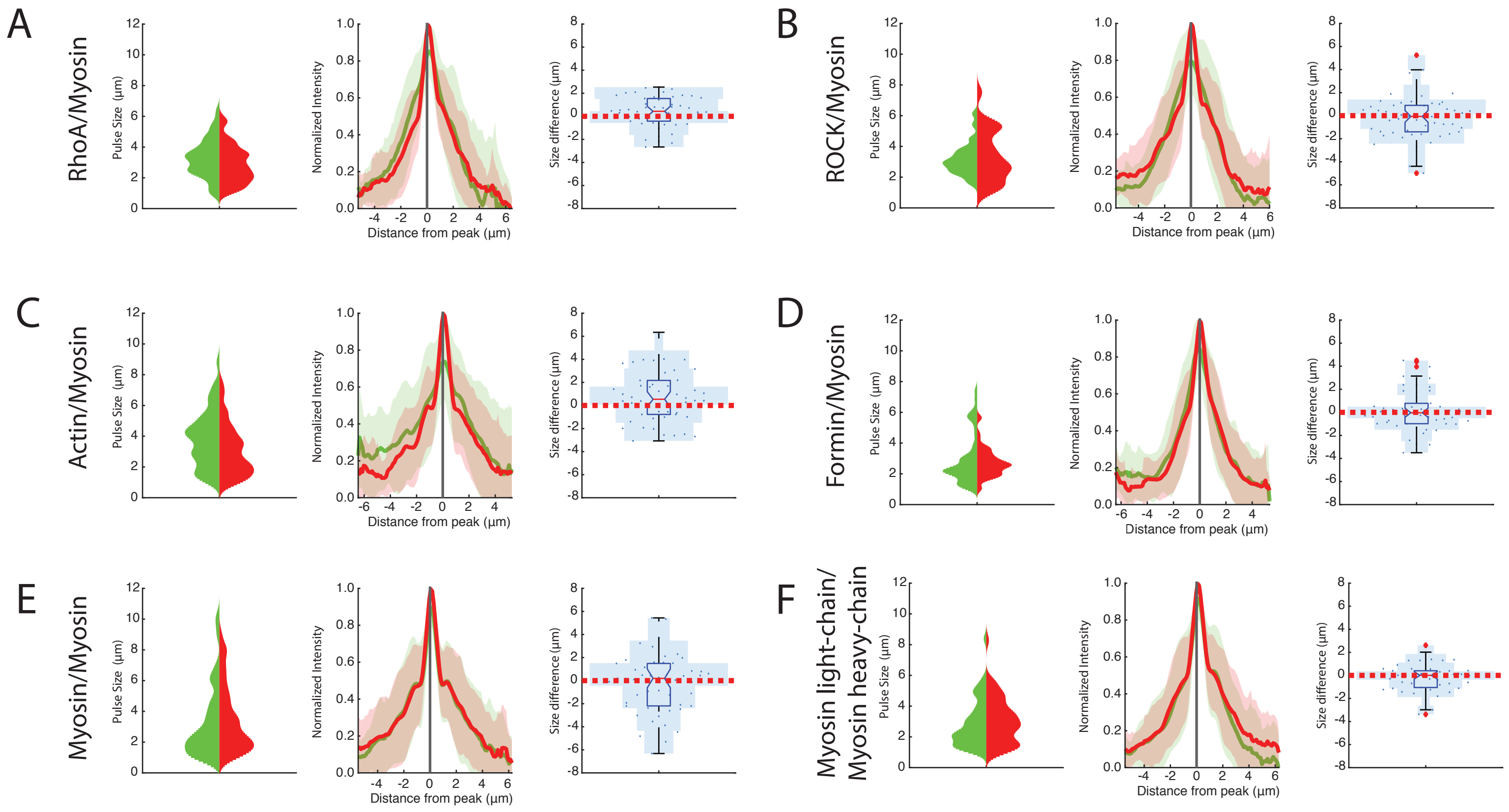

A
RhoA/Myosin
B
ROCK/Myosin
C
Actin/Myosin
D
Formin/Myosin
E
Myosin/Myosin
F
Myosin light-chain/
Myosin heavy-chain
Pulse Size (µm)
Normalized Intensity
Distance from peak (µm)
Size difference (µm)

**Supplemental figure 2. The size of Myosin pulse is smaller compared to the one of RhoA and Actin.** Left: distribution of pulse sizes, center: mean of the pulse size (solid line) with the std (shade), right: quantification of the difference in size (green – red). N(embryos) = 10, N(pulses) = 50. See statistical details in Supplementary Table S3. **(A-F)** Comparison in pulse sizes between in red Myosin (NMY-2) and in green RhoA (AHPH) (A), ROCK (LET-502) (B), Actin (UTR) (C), Formin (CYK-1) (D), the control Myosin (E) and Myosin Light-Chain (MLC-4) (F).

### Figure S3

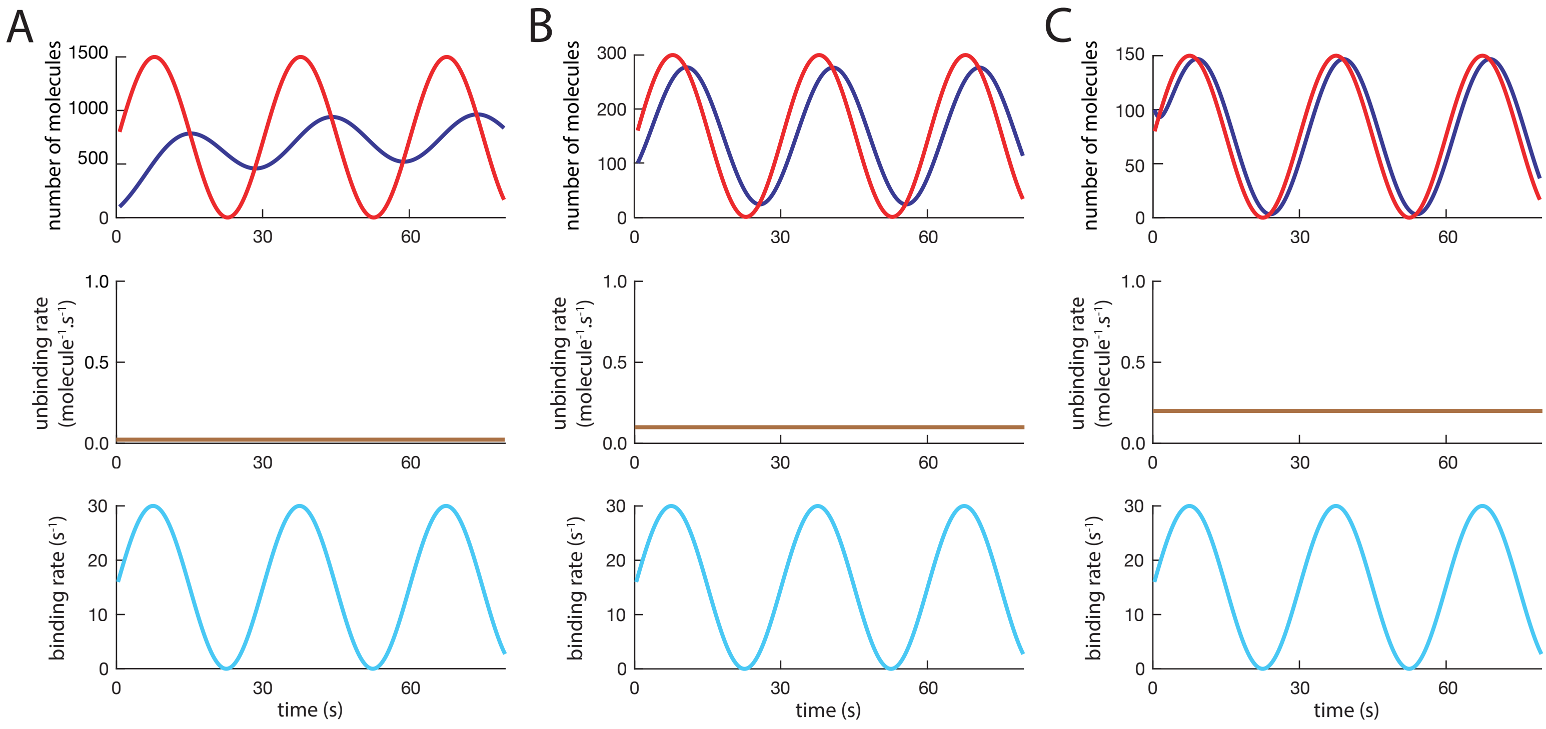

A
B
C
number of molecules
1500
1000
500
0
300
200
100
150
100
50
unbinding rate
(molecule$^{-1}$.s$^{-1}$)
1.0
0.5
0.0
binding rate (s$^{-1}$)
30
20
10
0
30
60
time (s)

**Supplemental figure 3. The arbitrary constant number used for the $K_{off}$ influences the delay between the simulated and the calculated number of particles. (A-C)** The simulations (blue line) shows that the system is more delayed and is further apart from the equilibrium state (red line: $K_{on}/k_{off}$) when the constant is lower like in (A) $k_{off}$ = 0.01 molecules$^{-1}$.s$^{-1}$, than it is when the $k_{off}$ is higher like in (B) $k_{off}$ = 0.1 molecules$^{-1}$.s$^{-1}$. When the $k_{off}$ is high like (C) $k_{off}$ = 0.2 molecules$^{-1}$.s$^{-1}$ then the simulation almost matches the equilibrium state and time delay is minimal.

### Figure S4

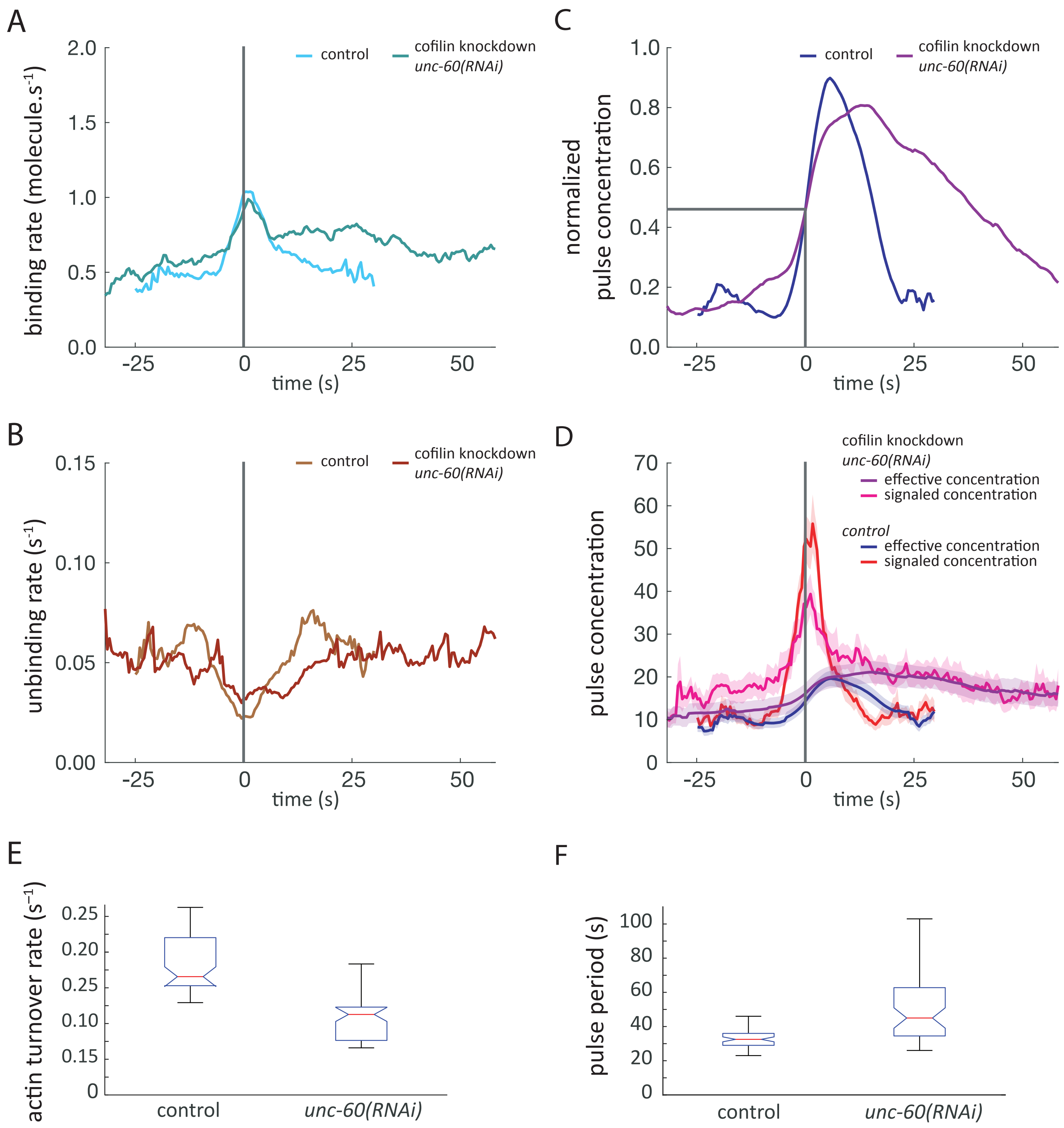
A
binding rate (molecule.s$^{-1}$)
2.0
1.5
1.0
0.5
0.0
-25
0
25
50
time (s)
control
cofilin knockdown
*unc-60(RNAi)*
B
unbinding rate (s$^{-1}$)
0.15
0.10
0.05
0.00
control
cofilin knockdown
*unc-60(RNAi)*
C
normalized pulse concentration
1.0
0.8
0.6
0.4
0.2
0.0
control
cofilin knockdown
*unc-60(RNAi)*
D
pulse concentration
70
60
50
40
30
20
10
0
cofilin knockdown
*unc-60(RNAi)*
effective concentration
signaled concentration
*control*
effective concentration
signaled concentration
E
actin turnover rate (s$^{-1}$)
0.25
0.20
0.25
0.10
0.15
0
control
*unc-60(RNAi)*
F
pulse period (s)
100
80
60
40
20
0
control
*unc-60(RNAi)*

**Supplemental figure 4. Cofilin depletion results in increased pulse period. (A-D)** Comparison between wild-type (dark blue) and Cofilin knockdown *unc-60(RNAi)* (light blue). Thin line: median, thick line: average. (A-C) Shade: std. **(A)** Binding rate ($K_{on}$) of Myosin over time, **(B)** unbinding rate ($k_{off}$), **(C)** Normalized pulse density. **(D)** Effective density, with control (dark blue) and *unc-60(RNAi)* (light blue) and signaled density ($K_{on}/k_{off}$), with control (red), *unc-60(RNAi)* (pink). Shade: sem. **(E)** Actin turnover rates from Actin::GFP single-molecule microscopy measurements in control and Cofilin (RNAi). (F) Pulse period measured from Actin::GFP TIRF microscopy movies in control and Cofilin (RNAi).

Table S1

| | n. of embryos | n. of pulses | t-test | delay median (s) | delay mean (s) | delay std |
|---|---|---|---|---|---|---|
| GFP::AHPH;NMY-2::mKate2 | 10 | 247 | 1.61E-102 | 4.3 | 4.6 | 2.0 |
| GFP::LET-502;NMY-2::mKate2 | 10 | 249 | 4.49E-73 | 4.6 | 4.6 | 2.8 |
| UTR::GFP;NMY-2::mKate2 | 10 | 249 | 1.52E-08 | 0.5 | 0.9 | 2.3 |
| CYK-1::GFP;NMY-2::mKate2 | 10 | 200 | 1.06E-66 | 4.6 | 4.8 | 2.6 |
| NMY-2::GFP;NMY-2::mKate2 | 10 | 200 | 8.67E-01 | 0.0 | 0.0 | 1.8 |
| MLC-4::GFP;NMY-2::mKate2 | 11 | 165 | 2.22E-02 | 0.5 | 0.3 | 1.6 |
| CAP-1::GFP;NMY-2::mKate2 | 10 | 200 | 2.51E-08 | 0.5 | 1.4 | 3.4 |
| MEL-11::GFP;NMY-2::mKate2 | 10 | 148 | 1.88E-04 | 1.5 | 2.2 | 6.7 |
| NMY-2::GFP SiMI (GFP RNAI) | 5 | 38 | | | | |
| NMY-2::GFP SiMI (unc-60 + GFP RNAi) | 8 | 39 | | | | |
| CYK-1::GFP SiMI | 8 | 56 | | | | |
| ACTIN::GFP SiMI | 6 | 55 | | | | |
| NMY-2::GFP SiMI (mel-11(it26) + GFP RNAi) | 9 | 54 | | | | |

Table S2

| | two-sample t-test |
|---|---|
| AHPH::GFP;NMY-2::mKate2 vs GFP::LET-502;NMY-2::mKate2 | 9.3E-01 |
| AHPH::GFP;NMY-2::mKate2 vs UTR::GFP;NMY-2::mKate2 | 1.2E-62 |
| GFP::LET-502;NMY-2::mKate2 vs UTR::GFP;NMY-2::mKate2 | 7.9E-48 |
| GFP::LET-502;NMY-2::mKate2 vs CYK-1::GFP;NMY-2::mKate2 | 5.3E-01 |
| UTR::GFP;NMY-2::mKate2 vs CYK-1::GFP;NMY-2::mKate2 | 1.2E-49 |
| UTR::GFP;NMY-2::mKate2 vs NMY-2::GFP;NMY-2::mKate2 | 2.8E-05 |
| MLC-4::GFP;NMY-2::mKate2 vs NMY-2::GFP;NMY-2::mKate2 | 1.4E-01 |
| CAP-1::GFP;NMY-2::mKate2 vs NMY-2::GFP;NMY-2::mKate2 | 6.4E-07 |
| UTR::GFP;NMY-2::mKate2 vs CAP-1::GFP;NMY-2::mKate2 | 5.8E-02 |

Table S3

| pulse size difference | one-sample t-test | median size Myosin (µm) | mean size Myosin (µm) | std size Myosin | median size GFP tagged protein (µm) | mean size GFP tagged protein (µm) | std size GFP tagged protein | median size difference (µm) | mean size difference (µm) | std size difference |
|---|---|---|---|---|---|---|---|---|---|---|
| RhoA vs Myosin Heavy-Chain | 0.04 | 2.70 | 2.89 | 1.23 | 3.27 | 3.28 | 1.26 | 0.43 | 0.39 | 1.31 |
| ROCK vs Myosin Heavy-Chain | 0.54 | 2.93 | 3.35 | 1.68 | 3.04 | 3.19 | 1.07 | -0.06 | -0.16 | 1.87 |
| Formin vs Myosin Heavy-Chain | 0.46 | 2.61 | 2.75 | 0.97 | 2.27 | 2.94 | 1.66 | -0.04 | 0.19 | 1.81 |
| Actin vs Myosin Heavy-Chain | 0.02 | 2.72 | 3.08 | 1.68 | 3.85 | 3.83 | 1.82 | 0.54 | 0.75 | 2.19 |
| Light Chain vs Myosin Heavy-C | 0.23 | 2.81 | 3.05 | 1.52 | 2.30 | 2.85 | 1.55 | 0.00 | -0.20 | 1.16 |
| Control : Myosin Heavy-Chain | 0.46 | 2.76 | 3.51 | 2.18 | 2.41 | 3.25 | 2.34 | 0.20 | -0.27 | 2.55 |

Table S4

| Strain name | genotype | source |
|---|---|---|
| EM264 | *nmy-2(cp52[nmy-2::mKate2 + LoxP unc-119(+) LoxP]) I; xsSi5[cb-unc-119 (+) GFP::ANI-1(AH+PH)] II; unc-119(ed3) III* | Michaux et al., 2018 |
| JH1541 | *unc-119(ed4) III; pJH7.03[pie-1p::GFP::actin::pie-1 3' UTR + unc-119(+)]* | Courtesy of G. Seydoux |
| LP229 | *nmy-2(cp52[nmy-2::mKate2 + LoxP unc-119(+) LoxP]) I; unc-119 (ed3) III* | Dickinson et al, 2017 |
| SWG282 | *gesIs008[cyk-1p::cyk-1::GFP::cyk-1UTR, unc-119+]* | Costache, Prigent Garcia |
| FBR175 | *nmy-2(cp52[nmy-2::mKate2 + unc-119(+)]) I; cyk-1(jme14[cyk-1::eGFP]) unc-119(ed3) III* | Costache, Prigent Garcia |
| FBR10 | *nmy-2(cp52[nmy-2::mKate2 + unc-119(+)]) I; xsSi3[cb-unc-119(+) pie-1::GFP::utrophin::pie-1 3' UTR] II; unc-119(ed3) III* | Tse et al., 2012 |
| ML2508 | *let-502(mc74[GFP::let-502]) I* | Bell et al., 2020 |
| FBR28 | *let-502(mc74[GFP::let-502]) nmy-2(cp52[nmy-2::mKate2 + LoxP unc-119(+) LoxP]) I; unc-119 (ed3) III* | this study |
| JJ1473 | *unc-119(ed3) III; zuIs45[nmy-2p::nmy-2::GFP + unc-119(+)] V* | Nance et al., 2003 |
| FBR189 | *nmy-2(cp52[nmy-2::mKate2 + LoxP unc-119(+) LoxP]) I; unc-119 (ed3) III; zuIs45 [nmy-2p::nmy-2::GFP + unc-119(+)] V* | this study |
| FBR96 | *mlc-4(jme4[mlc-4::eGFP + LoxP]) III* | this study |
| FBR119 | *nmy-2(cp52[nmy-2::mKate2 + LoxP unc-119(+) LoxP]) I; mlc-4(jme4[mlc-4::eGFP+loxP]) unc-119 (ed3) III* | this study |
| FBR157 | *mel-11(syb753[mel-11::GFP]) II* | this study |
| FBR227 | *nmy-2(cp52[nmy-2::mKate2 + LoxP unc-119(+) LoxP]) I; mel-11 (syb753[mel-11::GFP]) II; unc-119 (ed3) III* | this study |
| ML2519 | *cap-1(mc76[cap-1::GFP + unc-119(+)]) IV* | this study |
| FBR212 | *nmy-2(cp52[nmy-2::mKate2 + LoxP unc-119(+) LoxP]) I; cap-1(mc76[cap-1::GFP + unc-119(+)]) IV* | this study |
| SWG282 | *gesIs008[Pcyk-1::CYK-1::GFP::cyk-1UTR, unc-119+]* | Costache, Prigent Garcia |
| KK332 | *mel-11(it26) unc-4(e120) sqt-1(sc13)/mnC1 [dpy-10(e128) unc-52(e444)] II* | CGC |
| FBR236 | *mel-11(it26) unc-4(e120) sqt-1(sc13)/mnC1 [dpy-10(e128) unc-52(e444)] II; unc-119(ed3) III; zuIs45[nmy-2p::nmy-2::GFP + unc-119(+)] V* | this study |

### Analytical solution of myosin recruitment under a driving recruitment oscillatory signal

François B. ROBIN, Étienne PINARD

February 10, 2026

**Abstract**

Here we present the equations underlying a simplified model for myosin turnover under periodic pulses of an upstream signaling cascade that controls myosin cortical recruitment $k_{\text{on}}$. Simply, the model assumes pulsed periodic sinusoidal recruitment of myosin to the cortex from a large (infinite) cytoplasmic pool. We will first present the hypotheses that this model assumes, its basic equation, and solve it, reaching a form that makes explicit the attenuation of oscillation amplitude depending on pulse frequency (or angular frequency of recruitment oscillations, hereon $\omega$) and myosin turnover $k_{\text{off}}$.

#### 1 Assumptions underlying the model

In the model we propose, we make the following assumptions:

1. the cytoplasmic pool is well mixed,
2. the cytoplasmic pool is large compared to the cortical pool, that is, we can disregard the contribution of the myosin recruited to the cortex on the myosin available in the cytoplasm for binding,
3. there is a single cortical turn-over rate for myosin,
4. the myosin off-rate is not controlled by the upstream signaling cascade.

It should be noted that all of these assumptions are made also in experiments such as FRAP, FLIP, FDAP, ...

We consider the first-order linear ordinary differential equation

$$\frac{d[\text{Myo}]}{dt} = k_{\text{on}}^{\text{avg}} \sin^2(\omega t) - k_{\text{off}}[\text{Myo}], \tag{1}$$

where $k_{\text{on}}^{\text{avg}}$ is the average recruitment rate, $\omega$ is the angular frequency of oscillations, and $k_{\text{off}}$ is the dissociation rate constant.

#### 2 Trigonometric identity

Using the identity

$$\sin^2(\omega t) = \frac{1}{2}\left(1 - \cos(2\omega t)\right), \tag{2}$$

Eq. (1) can be rewritten as

$$\frac{d[\text{Myo}]}{dt} + k_{\text{off}}[\text{Myo}] = \frac{k_{\text{on}}^{\text{avg}}}{2} - \frac{k_{\text{on}}^{\text{avg}}}{2}\cos(2\omega t). \tag{3}$$

#### 3 Integrating factor

Equation (3) is linear. The integrating factor is

$$\mu(t) = e^{k_{\text{off}} t}. \tag{4}$$

Multiplying Eq. (3) by $\mu(t)$ gives

$$\frac{d}{dt}\left(e^{k_{\text{off}} t}[\text{Myo}]\right) = \frac{k_{\text{on}}^{\text{avg}}}{2}e^{k_{\text{off}} t} - \frac{k_{\text{on}}^{\text{avg}}}{2}e^{k_{\text{off}} t}\cos(2\omega t). \tag{5}$$

Equation (5) will be integrated piecewise. Using properties of exponential and trigonometric functions, we obtain the general solution.

$$[\text{Myo}](t) = \frac{k_{\text{on}}^{\text{avg}}}{2k_{\text{off}}} - \frac{k_{\text{on}}^{\text{avg}}}{2}\frac{k_{\text{off}}\cos(2\omega t) + 2\omega\sin(2\omega t)}{k_{\text{off}}^2 + 4\omega^2} + Ce^{-k_{\text{off}} t}. \tag{6}$$

The constant $C$ is determined by the initial condition.

#### 4 Periodic steady state

As $t \to \infty$, the transient term vanishes and the system approaches a periodic steady state.

Introducing an amplitude–phase representation,

$$k_{\text{off}}\cos(2\omega t) + 2\omega\sin(2\omega t) = \sqrt{k_{\text{off}}^2 + 4\omega^2}\cos(2\omega t - \phi), \tag{7}$$

with phase lag

$$\phi = \arctan\left(\frac{2\omega}{k_{\text{off}}}\right), \tag{8}$$

the steady-state solution can be written compactly as

$$[\text{Myo}]_{\text{ss}}(t) = \frac{k_{\text{on}}^{\text{avg}}}{2k_{\text{off}}} \cdot \left(1 - \frac{1}{\sqrt{1 + \frac{k_{\text{off}}^2}{4\omega^2}}} \cos\left(2\omega t - \phi\right)\right). \tag{9}$$

This form makes explicit the attenuation of oscillation amplitude depending on the angular frequency of recruitment oscillations $\omega$ and myosin turnover $k_{\text{off}}$:

$$\boxed{\textit{attenuation}: \sqrt{1 + \frac{k_{\text{off}}^2}{4\omega^2}}} \tag{10}$$
