## Supplementary material for "Kinetic Control of Out-Of-Equilibrium Dynamics in the RhoA Signaling Cascade Shapes Actomyosin Contractility": Movie Legends

**Movie S1. RhoA activation precedes Myosin II recruitment at the location of pulsed contraction.** Near-TIRF microscopy film of a 2-cell stage *C. elegans* embryo at interphase showing a proxy of RhoA activation in green (GFP::AHPH) and Myosin II in magenta (NMY-2::mKate2).

**Movie S2. ROCK recruitment precedes Myosin II recruitment at the location of pulsed contraction.** Near-TIRF microscopy film of a 2-cell stage *C. elegans* embryo at interphase showing ROCK in green (GFP::LET-502) and Myosin II in magenta (NMY-2::mKate2).

**Movie S3. Myosin Phosphatase accumulation does not correlate with Myosin II.** Near-TIRF microscopy film of a 2-cell stage *C. elegans* embryo at interphase showing Myosin Phosphatase in green (MEL-11::GFP) and Myosin II in magenta (NMY-2::mKate2).

**Movie S4. Myosin Heavy-Chain and Regulatory Light-Chain are recruited simultaneously at the location of pulsed contraction.** Near-TIRF microscopy film of a 2-cell stage *C. elegans* embryo at interphase showing Myosin Regulatory Light-Chain in green (MLC-4::GFP) and Myosin II Heavy-Chain in magenta (NMY-2::mKate2).

**Movie S5. Single-particle tracking of Myosin molecules during pulsed contractions.** Near-TIRF microscopy film of Myosin II (NMY-2::GFP) at single-molecule level at 2-cell stage at interphase. Top movie shows the tracking of molecules of the bottom movie.

**Movie S6. Formin recruitment precedes Myosin II recruitment at the pulsed contraction location.** Near-TIRF microscopy film of a 2-cell stage *C. elegans* embryo at interphase showing formin in green (CYK-1::GFP) and Myosin II in magenta (NMY-2::mKate2).

**Movie S7. F-actin accumulation precedes Myosin II during actomyosin pulsed contractions.** Near-TIRF microscopy film of a 2-cell stage *C. elegans* embryo showing a proxy for F-actin in green (UTR::GFP) and Myosin II in magenta (NMY-2::mKate2). Note that pulsed contractions can be observed in both cells (AB, left and P1, right) of the 2-cell stage embryo.

**Movie S8. Capping Protein recruitment can be seen briefly before Myosin II recruitment.** Near-TIRF microscopy film of a 2-cell stage *C. elegans* embryo at interphase showing Capping Protein in green (CAP-1::GFP) and Myosin II in magenta (NMY-2::mKate2).
